# Identification of circadian rhythms in *Nannochloropsis* species using bioluminescence reporter lines

**DOI:** 10.1101/550954

**Authors:** Eric Poliner, Cameron Cummings, Linsey Newton, Eva M. Farré

**Affiliations:** Cell and Molecular Biology Program, Michigan State University, East Lansing, MI 48824, USA; Department of Plant Biology, Michigan State University, East Lansing, MI 48824, USA

**Keywords:** circadian rhythms, stramenopiles, luciferase, transcription, cell division, bioluminescence reporter, *Nannochloropsis oceanica*, *Nannochloropsis salina*, *Nannochloropsis* genus

## Abstract

Circadian clocks allow organisms to predict environmental changes caused by the rotation of the Earth. Although circadian rhythms are widespread among different taxa, the core components of circadian oscillators are not conserved and differ between bacteria, plants, animals and fungi. Stramenopiles are a large group of organisms in which circadian rhythms have been only poorly characterized and no clock components have been identified. We have investigated cell division and molecular rhythms in *Nannochloropsis* species. In the four strains tested, cell division occurred principally during the night period under diel conditions, however, rhythms dampened within 2-3 days after transfer to constant light. We developed firefly luciferase reporters for long-term monitoring of *in vivo* transcriptional rhythms in two *Nannochlropsis* species, *N. oceanica* CCMP1779 and *N. salina* CCMP537. The reporter lines express free-running bioluminescence rhythms with periods of ~21-31 h that dampen within ~3-4 days under constant light. Using different entrainment regimes, we demonstrate that these rhythms are regulated by a circadian-type oscillator. In addition, the phase of free-running luminescence rhythms can be modulated pharmacologically using a CK1 ε/δ inhibitor, suggesting a role of this kinase in the *Nannochloropsis* clock. Together with the molecular and genomic tools available for *Nannochloropsis* species, these reporter lines represent an excellent system for future studies on the molecular mechanisms of stramenopile circadian oscillators.

**Significance statement:** Stramenopiles are a large and diverse line of eukaryotes in which circadian rhythms have been only poorly characterized and no clock components have been identified. We have developed bioluminescence reporter lines in *Nannochloropsis* species and provide evidence for the presence of a circadian oscillator in stramenopiles; these lines will serve as tools for future studies to uncover the molecular mechanisms of circadian oscillations in these species.

## Introduction

Circadian clocks generate physiological rhythms of periods of ~ 24 h in the absence of external cues. Circadian clocks are found in organisms that live exposed to daily light/dark cycles and allow them to predict and adapt to changes in their environment. In photosynthetic organisms, these clocks modulate photosynthetic capacity, growth, development, and responses to biotic and abiotic stimuli and have been shown to be necessary for optimal growth and survival (Dodd *et al.*, 2004, Gehan *et al.*, 2015, Ouyang *et al.*, 1998, Woelfle *et al.*, 2004, Yerushalmi *et al.*, 2011). However, some cyanobacteria and plant species apparently lack self-sustained rhythms (Gyllenstrand *et al.*, 2014, Holtzendorff *et al.*, 2008). Although evidence of rhythmic behavior has been detected in a large number of species in different taxa, a good understanding of the molecular components of circadian oscillators is only available for a few species of bacteria (Cohen and Golden 2015), archeoplastida (green algae and plants)(Linde *et al.*, 2017, Noordally and Millar 2015) and opisthokonts (animals and fungi) (Crane and Young 2014, Dunlap and Loros 2017). Little is known about the molecular mechanisms underlying the circadian clock in other eukaryotic lineages such as rhizaria, stramenopiles, or excavates.

In eukaryotes, circadian controlled transcription is regulated by a system of interlocked transcription-translation feedback loops. These circadian oscillators can be reset or entrained by environmental changes such as temperature and light, but are able to maintain similar periods across a wide range of ambient temperatures, a characteristic known as temperature compensation. In spite of the similarities among the structure of eukaryote circadian oscillators, their key components are not conserved among taxa. Studies on one or two model organisms in each taxon have driven our understanding of the mechanisms of these different circadian clocks (Bell-Pedersen *et al.*, 2005). The clocks of green algae and plants are characterized by the presence of pseudo-response regulators (PRRs) and the single MYB domain transcription factors (LHY), which were initially identified and characterized in *Arabidopsis thaliana*. The search for homologous genes have led the identification of clock components in other land plants and green algae, such as *Ostreococcus taurii* (Corellou *et al.*, 2009, Linde *et al.*, 2017). In a similar manner, the identification of the key components of animal clocks, which include basic helix-loop-helix PAS (Per/Arnt/Sim) proteins such us CLOCK, CYCLE and BMAL and animal type cryptochromes (Crane and Young 2014), was initially driven by work on fruit flies (Bargiello *et al.*, 1984, Zehring *et al.*, 1984). Comparative analyses then identified homologous clock genes in mammals (Tei *et al.*, 1997). Work on the fungus *Neurospora crassa* led to the discovery of the core components of fungal clocks, frequency (FRQ) and the LOV-domain containing light receptor white collar 1 (WC-1) (Crane and Young 2014). In this case, conservation of clock components has also been explored to investigate the role of the clock in other species (Hevia *et al.*, 2015, Lee *et al.*, 2018).

Stramenopiles or heterokonts are a diverse group of secondary endosymbionts, in which a previously non-photosynthetic eukaryote incorporated a red alga, and are phylogenetically distant from the other photosynthetic lineages such as Viridiplantae (green algae and plants) (Burki 2014). While stramenopiles include photosynthetic and non-photosynthetic groups, most stramenopiles are aquatic and comprise one of the most abundant groups found in phytoplankton (Rynearson and Palenik 2011). Transcriptome studies under light/dark cycles, have shown that a large proportion of transcripts is diel regulated in stramenopiles (Ashworth *et al.*, 2013, Chauton *et al.*, 2013, Gravot *et al.*, 2010, Poliner *et al.*, 2015, Smith *et al.*, 2016). Diel oscillations in sporulation and gene expression have been also been observed in the oomycete *Phytophtora infestans*, a non-photosynthetic stramenopile (Xiang and Judelson 2014). Self-sustained 24 h rhythms under constant environmental conditions have been measured in a few stramenopiles such as diatoms (Ragni and D’alcala 2007), brown algae (Schmid and Dring 1992, Schmid *et al.*, 1992) and *Nannochloropsis gaditana* (Braun *et al.*, 2014). However, other key characteristics of circadian oscillators such as entrainment by environmental signals or temperature compensation have not been investigated in stramenopiles.

Algae from the *Nannochloropsis* genus are a potential powerful model to characterize the circadian clock in stramenopiles. This genus of small non-motile unicellular algae includes fresh and marine species, although most studies have focused on salt water species due to their use as a source of fish food and omega-3 fatty acids, and their biotechnological potential. Diel rhythms in gene expression and cell division have been described in *N. oceanica* CCMP1779 and free running 24 h rhythms in chlorophyll content have been reported in *N. gaditana* CCMP1779 (Braun *et al.*, 2014). Several *Nannochloropsis* strains from different worldwide locations have been sequenced and current data suggests that *Nannochloropsis* species are haploid with genome sizes of ~30 Mb, containing from ~6,500 to ~12,000 genes (Corteggiani Carpinelli *et al.*, 2014, Radakovits *et al.*, 2012, Schwartz *et al.*, 2018, Vieler *et al.*, 2012, Wang *et al.*, 2014). Recent years have seen the development of a comprehensive set of molecular tools for working with these algae, such as homologous gene replacement, overexpression, gene silencing and CRISPR/Cas9 based genome editing (Kilian *et al.*, 2011, Poliner *et al.*, 2018a, Poliner *et al.*, 2018b, Poliner *et al.*, 2018c, Verruto *et al.*, 2018, Wang *et al.*, 2016, Wei *et al.*, 2017). In this study, we describe the development of a bioluminescence reporter system for the long-term *in vivo* measurement of gene expression in two *Nannochloropsis* species. We report a comprehensive analysis of entrainment and light and temperature dependence of circadian rhythms in *Nannochloropsis* genus, which form the base for future molecular characterization of the circadian oscillator in this stramenopile.

## Results

### *Nannochloropsis* species display oscillations in cell division under constant light conditions

*N. oceanica* CCMP1779 cultures grown under 12 h/12 h light/dark cycles synchronize their cell division, with DNA synthesis occurring during the second part of the light period and cytokinesis occurring during the night (Poliner *et al.*, 2015). To test if cell division is regulated by a circadian oscillator in *Nannochloropsis* species we measured cell numbers in cultures entrained under light/dark cycles and released into constant light (Fig. 1a). We analyzed four lines from two *Nannochloropsis* species originating from different latitudes (Figure S1). For our initial experiments, we chose a lower light intensity for the free running condition (40 μmol m^−2^ s^−1^) than the entrainment condition (100 μmol m^−2^ s^−^1) as has been used for experiments in the green marine algae *Ostreococcus tauri*. We observed cycling under light/dark for all four species. However, although all lines tested displayed a gated cell division during the second day under constant light conditions, two lines, CCMP537 and CCMP1778 did not display an oscillation during the first day in constant light. This transient effect could be due to the differences in light intensity under entrainment and free run conditions. We therefore analyzed growth in cells entrained and released at the same light intensity of 40 μmol m^−2^ s^−^1 (Fig. 1b). In this case, cell division in both *N. salina* CCMP537 and *N. oceanica* CCMP1779 was gated during the first day in constant light, but rhythms in the rate of cell division damped over the course of the experiment (Fig. 1c). The period estimate for *N. salina* CCMP537 using FFT-NLLS was 31 h, but no period estimation was possible for *N. salina* CCMP1779. Similar results were obtained with cells entrained and released at 100 μmol m^−2^ s^−1^ (Figure S2). These cell division rhythms in *Nannochloropsis* species are weaker than the tight synchronization observed in green algae (Goto and Johnson 1995, Moulager *et al.*, 2007) but similar to weak rhythms observed in the diatom *Phaeodactylum tricornutum* (Ragni and D’alcala 2007).

**Figure 1.**
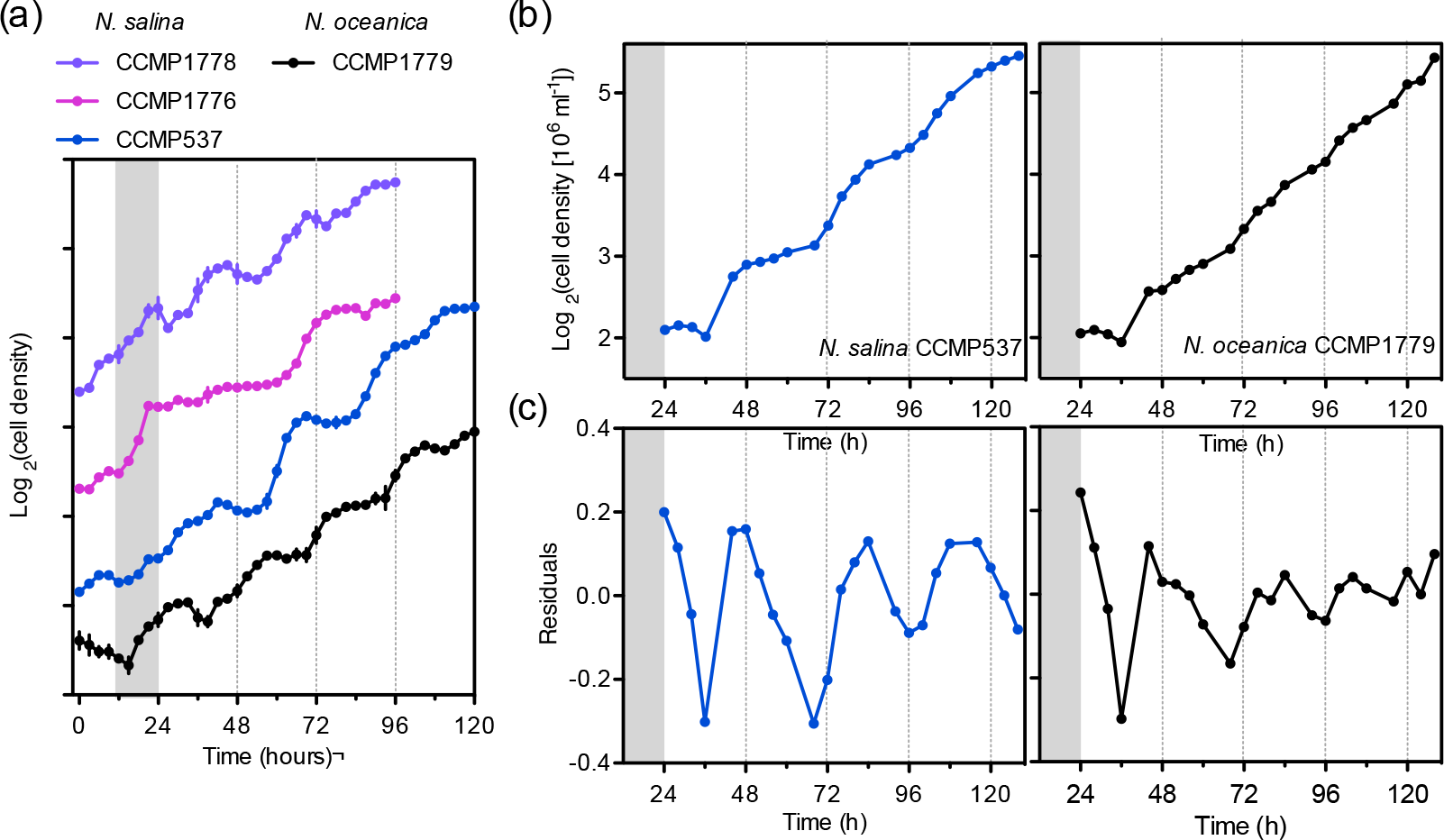
Cell division oscillations in *Nannochloropsis* species under light/dark and constant light conditions. **(a)** Cultures entrained under diel conditions (12 h light/12 h dark, 100 μmol m^−2^ s^−1^) and then moved to constant light (40 μmol m^−2^ s^−1^). Values are the average ± SEM (n=3 cultures) and traces are nudged to aid visualization. **(b)** Cultures grown under diel conditions (12 h light/12 h dark, 40 μmol m^−2^ s^−1^) for 10 days and then released to constant light (40 μmol m^−2^ s^−1^). Values are the average ± SEM (n=3, *N. salina*; n=2 *N. oceanica*). Grey shading indicates dark period. **(c)** Residuals after determining a linear regression of the plots shown in (b) to extract the oscillatory component.

### Development of luciferase reporter lines in *Nannochloropsis* species

In order to carry out a more detailed analysis of the rhythms observed in *Nannochloropsis* species, we developed bioluminescence reporter lines. We first tested three luciferase enzymes in the line *N. oceanica* CCMP1779, the Firefly luciferase (FLUC), Renilla luciferase (RLUC) and Nanoluciferase (NLUC). FLUC has been widely used as a circadian reporter in multiple systems (Welsh *et al.*, 2005, Welsh and Kay 2005), however, RLUC and the engineered NLUC have the advantage of being smaller proteins, and catalyze ATP-independent reactions (England *et al.*, 2016, Hall *et al.*, 2012). We have recently used NLUC to quantify protein expression in *N. oceanica* CCMP1779 (Poliner *et al.*, 2018c). The codon optimized luciferase coding regions were expressed under the control of the *CELLULOSE SYNTHASE* promoter (*CS*), a gene with strong oscillations in RNA levels under light/dark cycles (Poliner *et al.*, 2015). All *CS::LUCIFERASE* expressing lines displayed peaks of bioluminescence during the dark period in accordance to the expression of the endogenous *CS* gene (Fig. 2a)(Poliner *et al.*, 2015). Although RLUC and NLUC expressing lines showed high initial levels of luminescence both reporters displayed a strong decrease of signal during the course of an experiment. In the case of RLUC, this decrease is likely due to the chemical instability of its substrate, coelenterazine, in aqueous solution (Andreu *et al.*, 2010). The NLUC substrate, furimazine, was initially selected for its higher stability under animal tissue culture conditions (Hall *et al.*, 2012) and we tested whether instability under our culture conditions could explain the decrease in signal. Refeeding of substrate within a time course transiently increased the amplitude of oscillations of NLUC expressing lines but had little effect on FLUC luminescence, indicating that the NLUC substrate might be unstable under our experimental conditions (Figure S3). We also tested FLUC expression under the control of the *light harvesting complex 8* gene (*LHC8*) promoter in *N. oceanica* CCMP1779. The bioluminescence of this reporter peaked at dawn, which correlated to the morning expression of the *LHC8* gene in this strain (Fig. 2a)(Poliner *et al.*, 2015).

**Figure 2.**
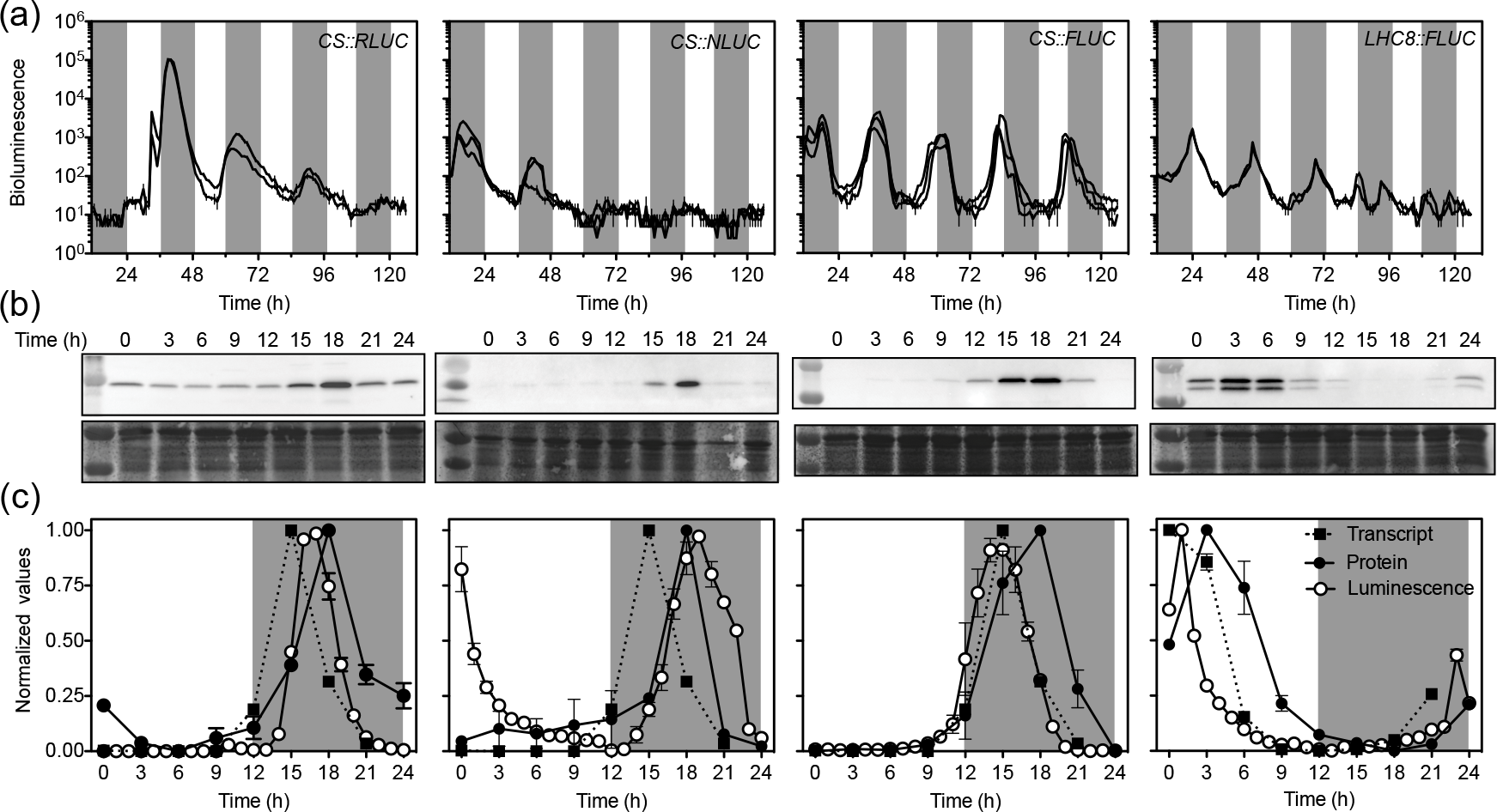
The development of bioluminescence reporters in *N. oceanica* CCMP1779. **(a)** *In vivo* luminescence oscillations of cells expressing *CS::RLUC*, *CS::NLUC, CS::FLUC* or *LHC8::FLUC* bioluminescence reporters under light/dark cycles. Dark grey shading indicates dark period. Luminescence was recorded every hour and the average bioluminescence per independent transgenic line ± SEM is shown (n=2 cultures). *CS*, *cellulose synthase* promoter; *LHC8*, *light harvesting complex 8* promoter; *RLUC*, *renilla luciferase*; *NLUC*, *nanoluciferase*; *FLUC*, firefly luciferase. **(b)** Representative western blots of the luciferase proteins from the transgenic reporter lines shown in **(a)** (top panel) and the blot stained with DB71 as loading control (bottom panel) over the course of one light/dark cycle. Light and dark periods are as indicated with grey shading in **c**. NLUC and FLUC contain a HA tag and were detected using an αE;-HA antibody. Renilla luciferase was detected using a specific antibody. **(c)** Quantitation of luciferase protein, and *in vivo* luminescence of one of the lines shown in (a), and the transcript abundance of the respective endogenous gene. Expression was normalized between 0 and 1 and the average ± SEM (n= 2 cultures) is shown. Transcript abundance data is from RNA-seq (Poliner *et al.*, 2015).

Under light/dark cycles, all the luciferase constructs tested led to oscillations in luciferase protein content that were delayed with respect to the RNA level of the corresponding gene, *CS* or *LHC8* (Fig. 2b, c). The doublet band in the *LHC8::FLUC* expressing lines is likely caused by an alternative translational start site within the *LHC8* upstream sequence. The *in vivo* FLUC bioluminescence correlated well with RNA levels of the *CS* gene (Fig. 2c). In contrast, the maximum RLUC and NLUC *in vivo* bioluminescence were delayed with respect to the *CS* RNA levels. These results indicate that *in vivo* firefly luciferase enzyme has a short catalytic activity half-life in *Nannochloropsis* species. In animal cells, firefly luciferase also responds faster than nanoluciferase (Hall *et al.*, 2012), which might be due to the product inhibition of the firefly luciferase enzyme (Leitao and Esteves da Silva 2010). Therefore, FLUC *in vivo* luminescence in the presence of substrate appears to reflect the rate of production of newly synthesized luciferase enzyme, which in our case, correlates with its RNA content, in a similar manner to what has been observed in higher plants (Farre and Kay 2007, Millar *et al.*, 1992).

Due to the robust activity of FLUC reporters in *N. oceanica* CCMP1779, we also developed luciferase reporter lines in *N. salina* CCMP537. *N. salina* strains display oscillations in cell division (Fig. 1) and the genome of *N. salina* CCMP537 has been sequenced (Wang *et al.*, 2014). In *N. salina* CCMP537, the FLUC reporter under the control of its endogenous *CS* promoter also maintained high amplitude oscillations under light/dark cycles with bioluminescence maxima during the dark period (Fig. 3). As in *N. oceanica* CCMP1779, *LHC8* driven FLUC *in vivo* bioluminescence in *N. salina* CCMP537 displayed a dawn peak of expression that weakened during the course of the experiment. The differences in the time of maximum expression between *CS* and *LHC8* driven FLUC in both *Nannochloropsis* species tested indicate that the oscillations of *in vivo* FLUC activity are not due to changes of endogenous ATP levels, a substrate of firefly luciferase, but reflect differences in transcriptional activity of these promoters during the light/dark cycle. We therefore, focused on *N. oceanica* CCMP1779 and *N. salina* CCMP537 *CS::FLUC* lines to further characterize the circadian rhythms of *Nannochloropsis* species. For brevity, these lines are described from now on as *N. oceanica* and *N. salina* in the text and figures.

**Figure 3.**
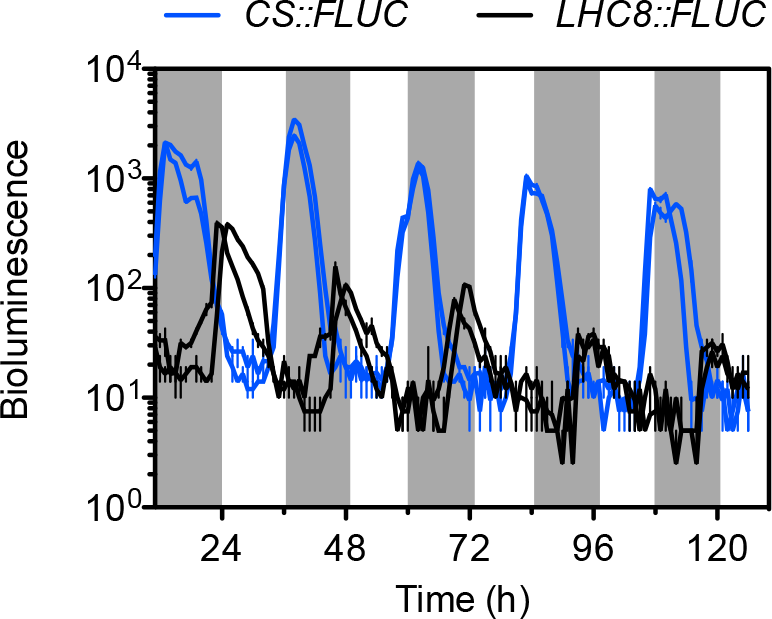
Bioluminescence oscillations of *N. salina* expressing *CS::FLUC* or *LHC8::FLUC* under light/dark cycles. Grey shading indicates dark period. Luminescence was recorded every hour and the average bioluminescence per independent transgenic line ± SEM is shown (n=2). *CS*, *cellulose synthase* promoter; *LHC8*, *light harvesting complex 8* promoter; *FLUC*, *firefly luciferase.*

### *Nannochloropsis* species display damping oscillations in gene expression under constant light conditions

Under light/dark cycles, both *CS::FLUC* and *LHC8:FLUC* reporters displayed anticipatory behavior under light/dark cycles indicating the presence of a endogenous oscillator (Fig. 2, 3). To test if *Nannochloropsis* species maintain transcriptional rhythms under constant light we measured *in vivo* bioluminescence in *N. salina* and *N. oceanica* expressing the *CS::FLUC* reporter, which exhibited higher amplitude rhythms (Fig. 4). Since rhythms in the marine green algae *Ostreococcus tauri* are lost at moderate to high light intensities we first tested luminescence rhythms under different intensities of white light (Moulager *et al.*, 2007).

**Figure 4.**
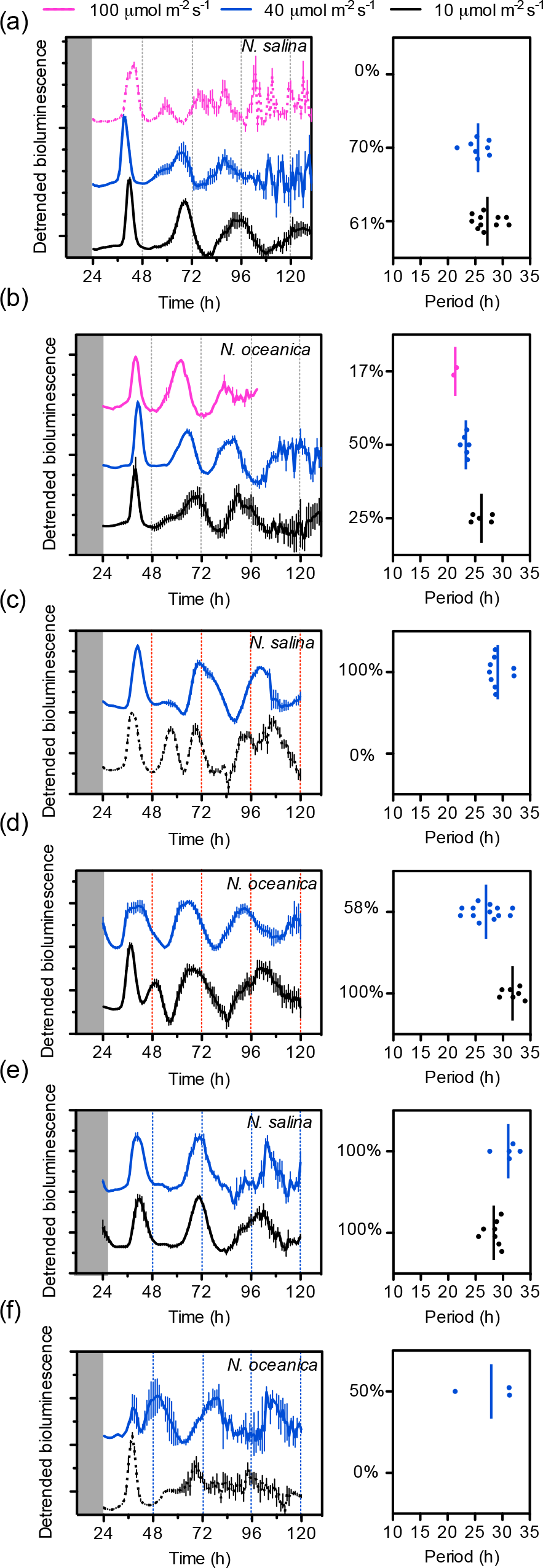
*In vivo* bioluminescence oscillations of *CS::FLUC* expressing lines from two *Nannochloropsis* species in constant light. Cultures were entrained under cycles of 12 h light/12 h dark (200 μmol m^−2^ s^−1^) and 22°C and released under either constant white **(a,b**), red **(c,d**) or blue light **(e,f**) at the indicated intensities (100 μmol m^−2^ s^−1^, magenta; 40 μmol m^−2^ s^−1^, blue; 10 μmol m^−2^ s^−1^, black). Right panels represent the average of detrended bioluminescence traces ± SEM (*N. salina* n=4-18 cultures, two independent lines; *N. oceanica* n=6-20, three independent lines). Only rhythmic traces are shown, with the exception of conditions that resulted in no rhythmic traces, for which the average trace is shown as a dotted line. Percent values indicate percent rhythmic traces. Traces are nudged to aid visualization. Left panels represent the period length estimate by FFT-NLLS on Biodare 2 using data from the respective right panel, only period lengths of traces considered rhythmic are shown, line indicates average.

Bioluminescence traces were detrended and the periods estimated using Biodare 2 (Moore *et al.*, 2014)(Figure S4). After entrainment under light/dark cycles, *N. salina* maintained rhythms for 2-3 cycles in constant white light conditions. The length of its free running period was ~27 h under 10 μmol m^−2^ s^−1^ and ~26 h under 40 μmol m^−2^ s^−1^, but under 100 μmol m^−2^ s^−1^ of white light no rhythmic cultures were observed. In contrast, *N. oceanica* displayed weaker rhythms than in *N. salina* under constant white light, with fewer rhythmic cultures (Fig. 4). In particular, under 10 μmol m^−2^ s^−1^ white light, only 25% of the cultures were rhythmic based on our rhythmic criteria (see Methods). Under weak red light, *N. salina* displayed erratic oscillations and no rhythmic cultures, however, better rhythms were detected under 40 μmol m^−2^ s^−1^ red light with an average period of 29 h. In contrast, *N. oceanica*’s rhythms were weaker under blue light than under red light. In the dark under autotrophic conditions, firefly luciferase reporter activity in both *Nannochloropsis* species was strongly decreased, as is the case with the marine unicellular algae *Ostreococcus taurii* (Figure S5) (O’Neill *et al.*, 2011).

### *In vivo* luminescence rhythms are temperature compensated and are maintained under short light/dark cycles

One of the key characteristics of circadian oscillators is their ability to maintain similar circadian periods under a wide range of temperatures, which is termed temperature compensation. We investigated *CS::FLUC* driven luminescence under low white light and temperatures ranging from 17°C to 28°C (Fig. 5a). *N. salina* displayed oscillations in all temperatures tested; all cultures were rhythmic under 19°C and 22° C, but only ~60% were rhythmic at the higher temperatures tested. In contrast, the rhythms were weak for *N. oceanica* at all temperatures tested and cyclic cultures were observed only under 19°C, 22°C and 25°C (Figure S6). In *N. salina* the estimated period lengths showed great variability, ranging from 23 to 31 h (Fig. 5b). We calculated the Q10, the factor by which the rate of a reaction varies in response to a 10 °C change, to quantify the degree of temperature compensation of *N. salina* bioluminescence rhythms. Using all the available data we estimated the temperature compensation for transcriptional rhythms in *N. salina* to have a Q10 of ~ 1.1, which lies within the range of other characterized circadian oscillations in plants(Kusakina *et al.*, 2014) and algae (Anderson *et al.*, 1985)(Fig. 5b).

**Figure 5.**
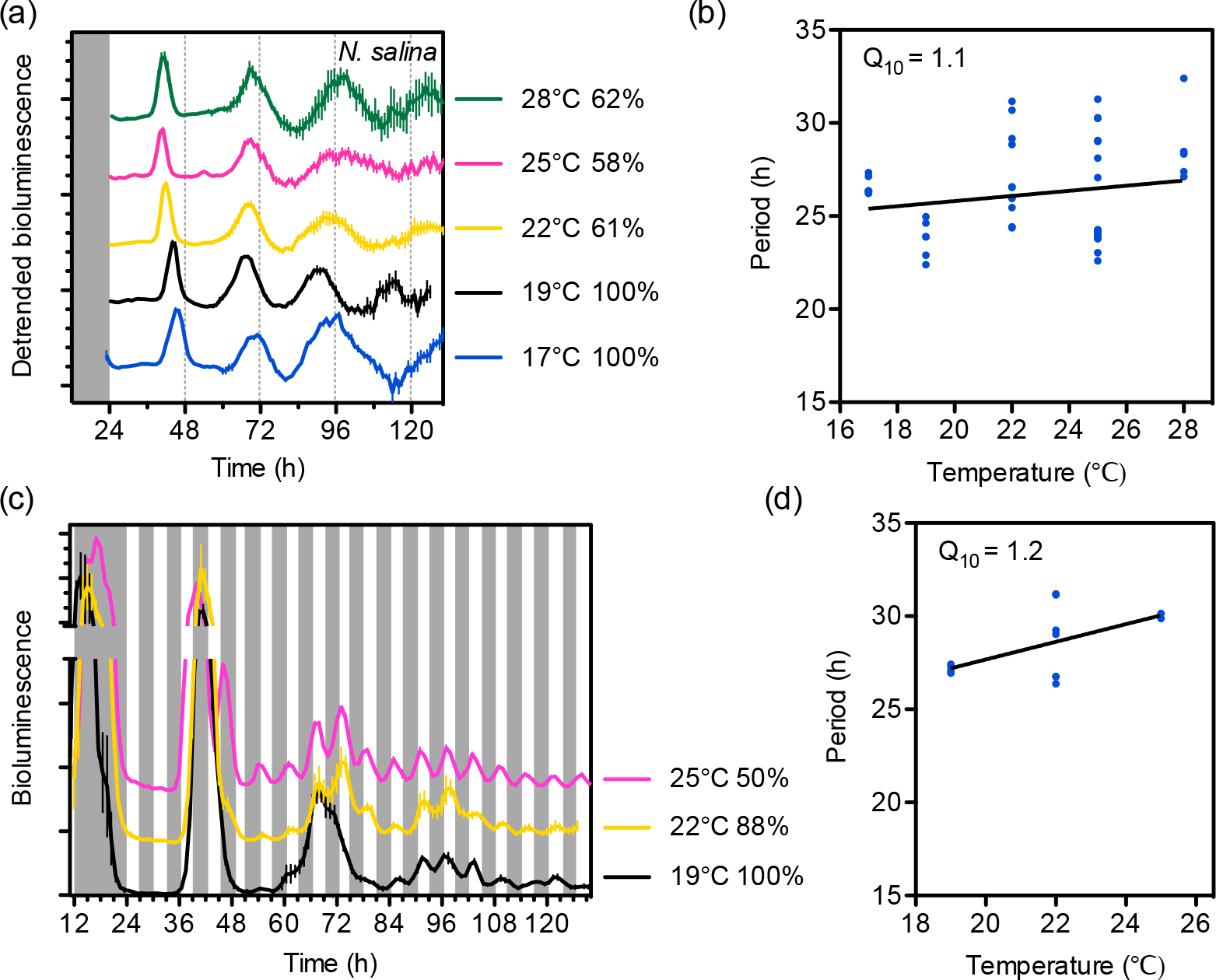
Rhythms of *N. salina CS::FLUC* expression under different temperatures. Cultures were entrained under cycles of 12 h light/12 h dark (200 μmol m^−2^ s^−1^) and 22°C. **(a)** *In vivo* bioluminescence under constant white light conditions (10 μmol m^−2^ s^−1^). Cultures were switched to 17°C or 19°C at time 12 h, and to 25°C or 28°C at time 24 h. The average detrended bioluminescence of rhythmic cultures ± SEM is shown (*N. salina* n=2-15, two independent lines; *N. oceanica* n=6-18, three independent transgenic lines). Traces are nudged to aid visualization. **(b)** Period lengths estimated by FFT-NLLS on Biodare 2 using data from *N. salina* shown in **(a)**. Only period lengths of traces considered rhythmic are shown. **(c)** *In vivo* bioluminescence of *CS::FLUC* in *N. salina* under T6 cycles. Cultures were transferred to cycles of 3 h white light (100 μmol m^−2^ s^−1^) and 3 h of dark at time 24 h. Grey shadings indicate dark periods. Cultures were switched to different temperatures as described for **(a)**. The average bioluminescence per rhythmic culture ± SEM of rhythmic traces is shown (n=2-7). Percent values indicate the percent of rhythmic traces. **(d)** Period length of traces shown in **c** estimated by FFT-NLLS on Biodare 2. Q10, factor by which the rate of a reaction varies in response to a 10°C change in temperature calculated using the slopes from **(b)** and **(d)** respectively.

Environmental factors that entrain circadian oscillators (zeitgebers), such as light, can lead to masking of the measured physiological rhythm. For example, light inhibits locomotor activity in mice and therefore mice activity is arrhythmic under constant light conditions (Ohta *et al.*, 2005). A similar effect could be involved in the damping rhythms of transcriptional luciferase reporters under constant light conditions in *Nannochloropsis* species. Since circadian oscillators change the time-dependent sensitivity towards the entrainment signal, in this case light, short light/dark cycles can reveal free-running rhythms, a process called frequency-demultiplication, in organisms in which masking occurs (Aschoff 1999). Therefore, when the endogenous period is a multiple of the frequency of the light/dark cycle, organisms with a circadian oscillator maintain their entrainment to the 24 h cycle and do not entrain to the new shorter cycles. To further test whether a circadian oscillator is present in *Nannochloropsis* species, we measured rhythms under short cycles of 3 h light and 3 h darkness (period = T= 6 h). Under T6 cycles both *Nannochloropsis* species revealed no ultradian oscillations during the first day but small amplitude 6 h oscillations were detectable the second day onwards, such that the *CS::FLUC* reporter was induced in the dark and repressed in the light (Fig. 5c, Figure S7a). At 22° C *N. oceanica* and *N. salina* cells maintained bioluminescence oscillations for three days with average periods of 23.8 h and 28.6 h respectively (Fig. 5b, Figure S7c). As observed for the bioluminescence rhythms under constant light, the circadian rhythms in *N. salina* were more robust than the oscillations from *N. oceanica*, and 88% *N. salina CS::FLUC* cultures scored as rhythmic in these experiments in contrast to 28% for *N. oceanica*. Similar period lengths were observed in *N. salina* cells expressing the *LCH8::FLUC* reporter although the amplitude of these oscillations was much reduced and the *LHC* reporters were sensitive to the short light/dark cycles throughout the treatment (Figure S7b,c).

We also quantified the circadian period at different constant temperatures in *N. salina* under T6 cycles (Fig. 5c,d). Rhythmic cultures were detected at all three temperatures tested (Fig. 5c, d). Cultures at 19°C lacked ultradian oscillations during both the first and second day in T6 conditions indicating a more robust circadian regulation than at higher temperatures. *N. salina CS::FLUC* rhythms under T6 had a Q10 of 1.2, similar to the rhythms under constant light conditions (Fig. 5b,c).

### Luminescence rhythms in *Nannochloropsis* species respond to changes by light cues

To determine if the *in vivo* luminescence rhythms could be reset by light we first carried out a simulated jet lag experiment, in which *N. salina CS::FLUC* cells were entrained under light/dark conditions and then treated with a night extension of 6 h (Fig. 6a). This treatment led to a phase advance of 1-2 h in the first day after the shift; the *N. salina* cultures were able to entrain to the new phase by the second day. The small phase shift and the delay in resetting indicates the presence of an endogenous oscillator, such that the oscillations in luminescence are not only driven by the light/dark cycles.

**Figure 6.**
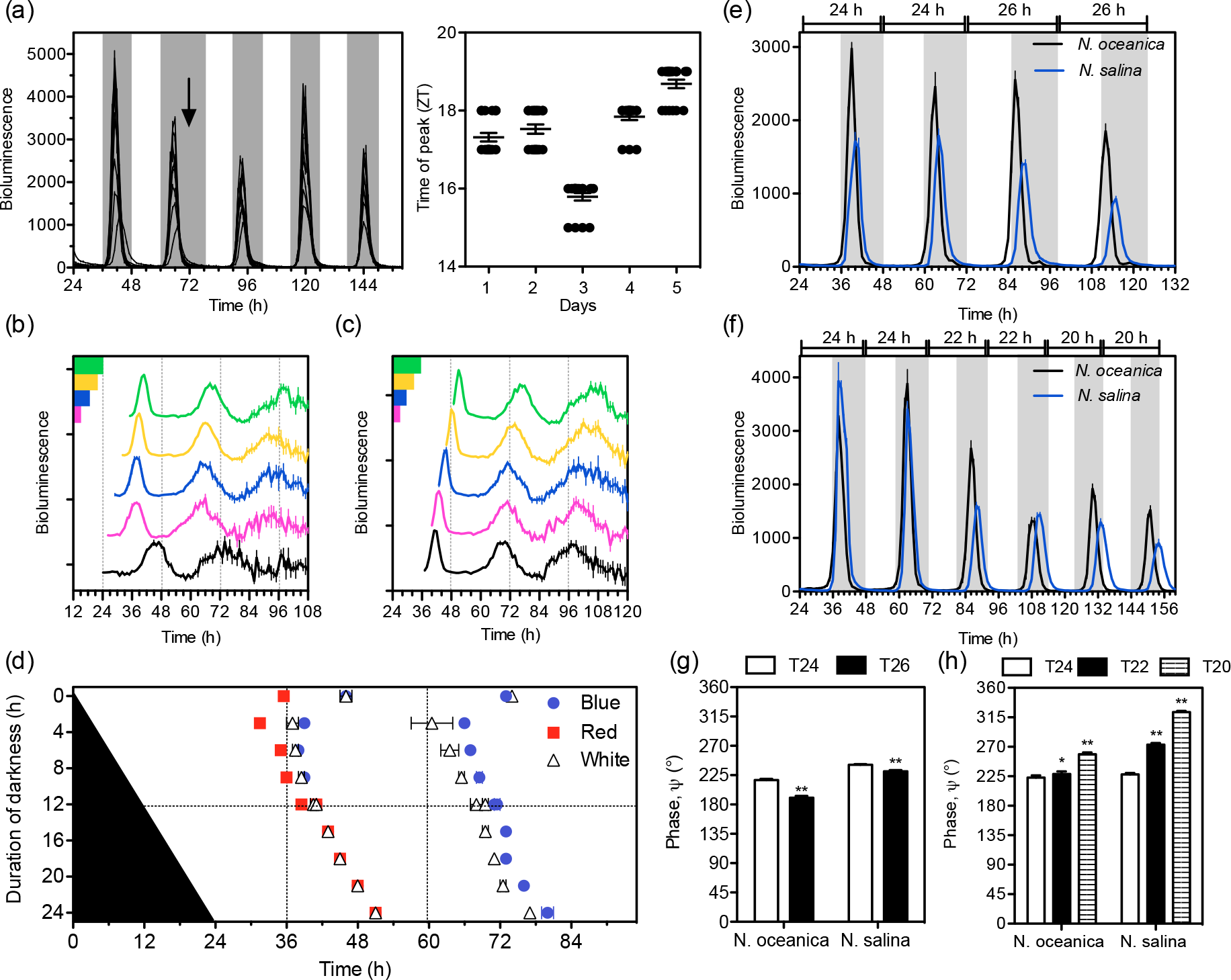
Light entrainment in *Nannochloropsis* species using *CS::FLUC* expressing lines. **(a)** Experimental jet-lag recovery in *N. salina*. Left panel, *in vivo* bioluminescence, traces from one representative experiment with cultures from two independent transgenic lines are shown. Grey shading indicates dark period. Cells were entrained under cycles of 12 h light/12 h dark and 22°C before the start of the experiment. At time 72 h (arrow) the night was extended by 6 h. Right panel, time of maximum luminescence with respect to last dawn (ZT in h) before (day 1 and 2) and after (days 3-4) the experimental jet-lag (average ± SEM; n= 19). **(b-d)** Phase response curve of *CS::FLUC* rhythms *in N. salina*. After entrainment under cycles of 12 h light/12 h dark and 22°C for 7 days, cultures were exposed to one dark period of variable length before being transferred to constant light (10 μmol m^−2^ s^−1^). Representative experiment using white light and extended nights **(b)** or short nights **(c)** (average ± SEM, n = 4, one transgenic line). For **(b)** the length of the dark periods were 12 h (black), 15 h (magenta), 18 h (blue), 21 h (yellow) or 24 h (green); and for **(c)**, 0 h (black), 3 h (magenta), 6 h (blue), 9 h (yellow) or 12 h (green). **(d)** Phase of peak luminescence (x-axis) versus the duration of the dark period (y-axis) after transfer to either white, blue or red light (average ± range of two independent transgenic lines). **(e-f)** *In vivo* luminescence under different T-cycles, with 30 μmol m^−2^ s^−1^ white light (average ± SEM, n = 16, two transgenic lines). **(g-h)** Phase angle of bioluminescence oscillations shown in (e) and (f), using traces from the second day in the respective T-cycle (average ± SEM, n = 16). Significant difference to T24 (*) p<0.05, (**) p<0.001(unpaired *t*-test).

To further investigate light entrainment in *N. salina* we carried out phase shift experiments under different light qualities. *N. salina CS::FLUC* cultures were grown under white 12h light/12 h dark cycles, treated with dark periods of different durations and then transferred to constant white, red or blue light conditions (Fig. 6b, c; Figure S8). As observed in the previous jet-lag experiments an extension of the dark period led to a delay in phase that correlated with the length of the dark extension (Fig. 6c,d, Figure S8). Dark periods shorter than 12 h led to phase advances of 2-3 h, which were shorter than the observed delays (Fig. 6b,d, Figure S8). These delays and advances were similar between blue and white light treatments; however, phase advances were slightly stronger under red light. In the absence of a dark period, the phase was delayed under blue and white light but not under red light. After the phase shifts, the new phase was maintained during the second day under free running conditions indicating a change in the phase of a circadian oscillator and not a direct light effect on the reporter expression (Fig. 6d). Under red light conditions rhythms dampened quickly (Figure S8b), making it difficult to determine the phase of the second peak of luminescence. These experiments further indicate the presence of a circadian oscillator in the genus *Nannochloropsis*.

Another characteristic of circadian oscillators is that they determine the phase relationship or phase angle of gene expression within a 24 h light/dark cycle (T24). Circadian control leads to phase advances when the cycles are longer than 24 h and two phase delays when the cycles are shorter than 24 h (Johnson *et al.*, 2003). We determined the phase angle, the relative time of peak expression of the *CS::FLUC* reporter within one cycle, in *N. salina* and *N. oceanica* under T20, T22, T24 and T26. Both species displayed stable entrainment under these conditions (Fig. 6e and f), however, the relative time of expression of the reporter within the light/dark cycles was advanced under T26 and delayed under T20 and T22 (Fig. 6g and h) supporting the presence of an endogenous oscillator.

### Entrainment of luminescence rhythms by temperature

In most organisms, temperature, in addition to light, plays an important role in resetting the circadian clock (Rensing and Ruoff 2002). We investigated entrainment of *CS::FLUC* oscillations in *Nannochloropsis* species by growing cultures under temperature cycles of 12 h 27°C and 12 h 17°C under constant light conditions. Temperature cycles were able to maintain robust high amplitude oscillations in both *N. salina* and *N. oceanica* (Fig. 7). The peak of *CS::FLUC* expression in both species occurred at the end of the warm period, an earlier phase than light entrainment. This phase relationship was maintained when cultures were transferred to constant temperature and light conditions. As observed for light entrained cultures, only a few *N. oceanica* cultures were rhythmic after temperature entrainment (Figure 7). For *N. salina*, the estimated average period of the luminescence rhythm after temperature entrainment, 28.8 h, was similar to the period after light entrainment.

**Figure 7.**
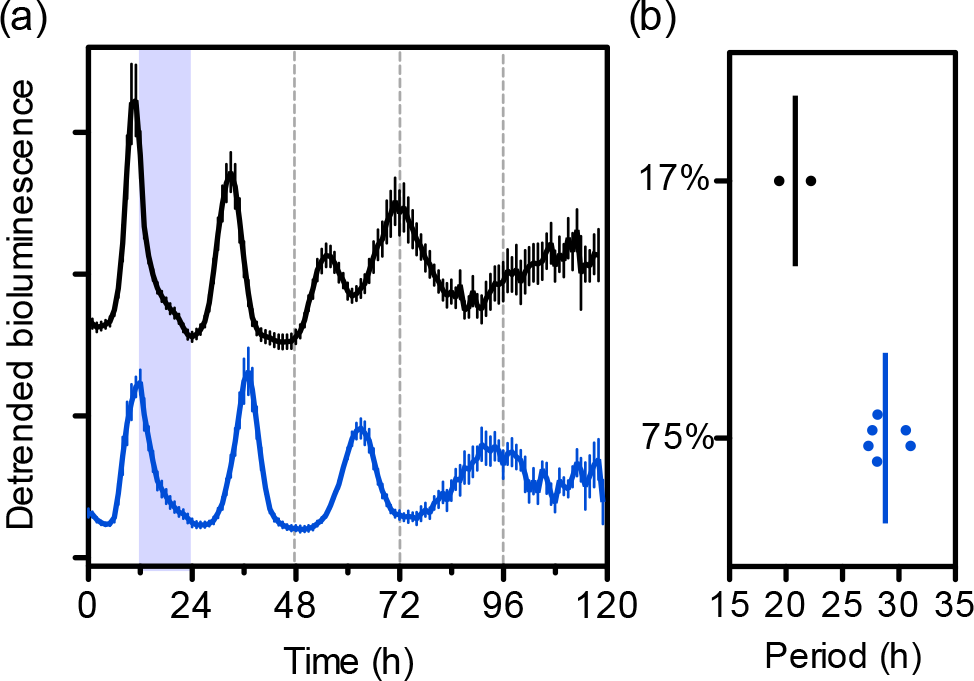
Temperature entrainment in *Nannochloropsis* species. **(a)** *In vivo* bioluminescence rhythms of *CS::FLUC* in *N. oceanica* (black) and *N. salina* (blue). Cultures were entrained under cycles of 12 h 17°C /12 h 27°C and 200 μmol m^−2^ s^−1^ white light. Bioluminescence was recorded for one warm/cool cycle (40 μmol m^−2^ s^−1^) and at time 24 h cells were transferred to constant temperature (22°C) and constant light (40 μmol m^−2^ s^−1^). The detrended bioluminescence ± SEM of cycling lines (*N. salina* n=6, two independent lines; *N. oceanica* n=3, three independent lines); purple shading indicates cool period, dotted lines indicate subjective cool to warm transition. Amplitude and baseline detrending was implemented on Biodare 2. **(b)** Period length of cycling cultures estimated using FFT-NLLS on Biodare 2. Percent cycling cultures are indicated.

### Potential clock components in the *Nannochloropsis* genomes

No circadian clock component has been identified in stramenopiles, however, *Nannochloropsis* genomes encode for genes with some similarities to plant and animal clock genes (Vieler *et al.*, 2012). Proteins containing plant-specific CCT domains are involved in circadian and photoperiod control in land plants and green algae (Farre and Liu 2013, Serrano *et al.*, 2009). We identified three CCT containing proteins annotated in the *N. oceanica* and the *N. gaditana* genomes and one in the *N. salina* genome (Figure S9). The CCT domains across stramenopiles display higher similarity to each other than to CCT domains outside that group. The expression of two of the CCT *N. oceanica* genes cycle under light/dark conditions (Figure S10a) (Poliner *et al.*, 2015). None of the currently annotated stramenopile CCT domain proteins appear to contain either response-regulator or BBOX domains, which are characteristic in green algae and land plant proteins with circadian or photoperiod function, and therefore, their potential role in the clock remains unclear.

*Nannochloropsis* species contain an animal cryptochrome like protein, homologe to the diatom CPF1 (cryptochrome photolyase family) (Vieler *et al.*, 2012) (Figure S11). Diatom CPF1 has been shown to be able to inhibit the transcriptional activity of the mouse clock proteins BMAL and CLOCK in a heterologous system, and has been proposed to be involved in circadian regulation (Coesel *et al.*, 2009). BMAL1 and CLOCK are part of the bHLH-PAS protein family, whose members are involved in circadian regulation in animals (Crane and Young 2014). bHLH-PAS proteins are not found in green algae or land plants but have been identified in red algae and stramenopile genomes including *Nannochloropsis* species (Thiriet-Rupert *et al.*, 2016, Vieler *et al.*, 2012)(Figure S12). Three of the four *N. oceanica* CCMP1779 bHLH-PAS proteins cycle under light/dark conditions (Figure S10b). In stramenopiles, these proteins contain only one PAS domain instead of the two PAS domains found in animal proteins (Figure S12a) and therefore it has been suggested that the stramenopile proteins are an example of convergent evolution (Thiriet-Rupert *et al.*, 2016).

Casein kinase 1 (CK1) is a conserved component of circadian systems in eukaryotes and the pharmacological inhibition of CK1 affects the circadian period in mammals (Badura *et al.*, 2007, Eide *et al.*, 2005, Walton *et al.*, 2009), *Drosophila melanogaster* (Meng *et al.*, 2010), *Neurospora crassa* (Querfurth *et al.*, 2011), *Ostreococcus tauri* (van Ooijen *et al.*, 2013) and *Caenorhabditis elegans* (Goya *et al.*, 2016). *N. oceanica* contains a CK1 gene with 72% identity with mouse casein kinase 1 ε (NannoCCMP1779|10930); this gene cycles under light/dark conditions with a peak at dawn (Figure S13a). To further test a possible involvement of a circadian oscillator in *Nannochloropsis* species, we investigated the effect of the CK1 ε/δ inhibitor PF-670462 on bioluminescence oscillations. In *N. salina*, PF-67462 led to phase delays of *CS::FLUC* expression in a concentration dependent manner (Fig. 8), although the free running period was only reduced in two of the tested concentrations (Figure S13b). PF-670462 treatment leads to a lengthening of the period in several organisms and causes phase delays in mammals (Badura *et al.*, 2007, van Ooijen *et al.*, 2013, Walton *et al.*, 2009). These results suggest that phosphorylation plays a key role in circadian control in *N. salina*.

**Figure 8.**
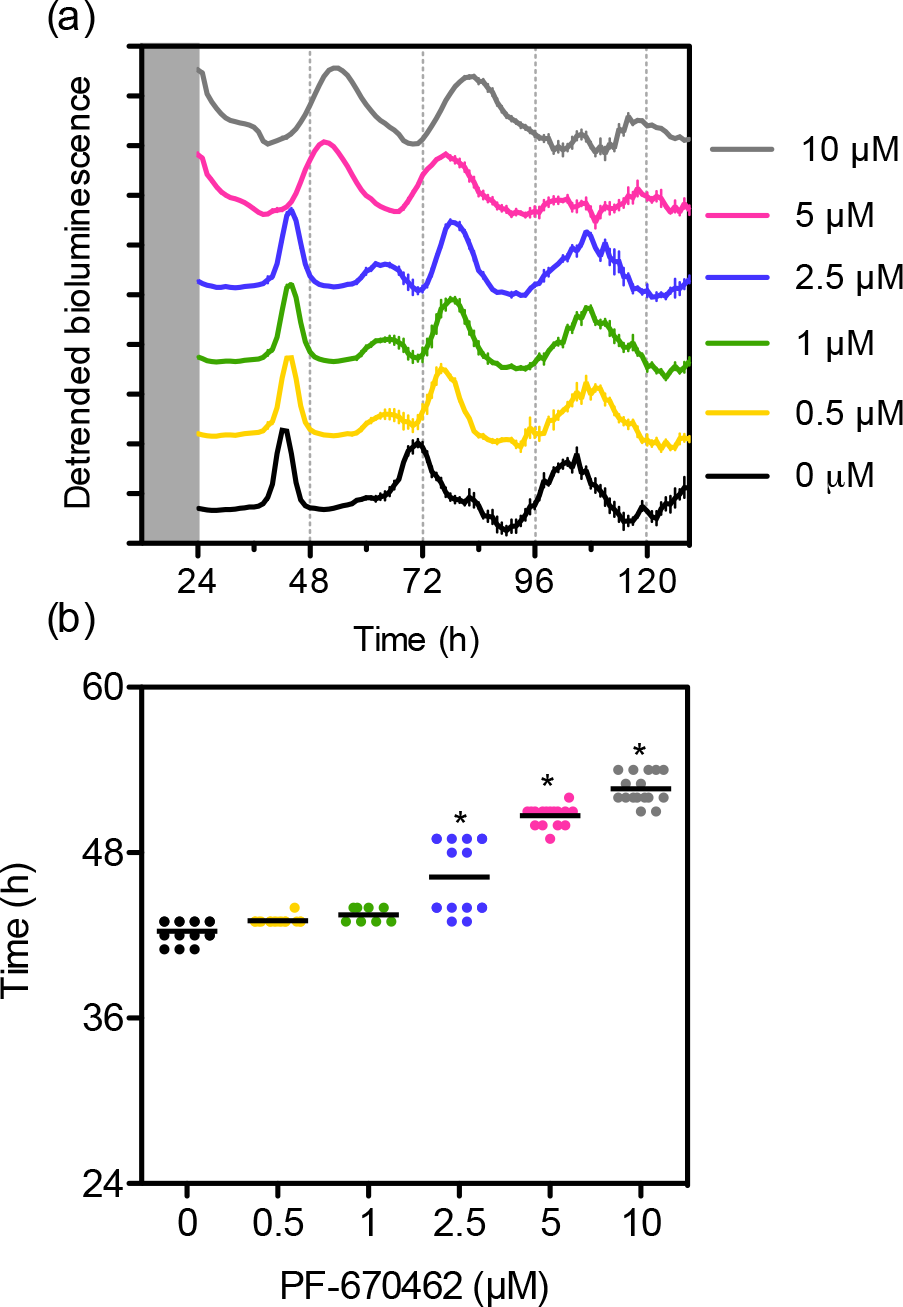
Modulation of *CS::FLUC* rhythms in *N. salina* by a CK1ε/δ inhibitor. **(a)** *In vivo* bioluminescence rhythms of *N. salina CS::FLUC* expressing lines. The average ± SEM from rhythmic traces is shown (n =3-8, two independent lines). Cultures were entrained under cycles of 12 h light/12 h dark (200 μmol m^−2^ s^−1^) and released to constant light (10 μmol m^−2^ s^−1^) at time 24 h. Cells were treated with PF-670462 at the indicated concentrations at time 12 h. Traces from one representative experiment are shown. Dark grey shading indicates dark period, light grey shading indicates subjective dark. **b** Time of first peak of *CS::FLUC* luminescence (n=8-16) from two independent experiments. * Indicate a significant difference with the vehicle control (one-way ANOVA with Dunnett’s post hoc test, α=0.05).

## Discussion

Circadian studies in stramenopiles have been limited to the measurement of physiological outputs such as cell division, pigment content or photosynthesis rates that are difficult to apply for high throughput studies (Makarov *et al.*, 1995, Ragni and D’alcala 2007). In this work, we describe oscillations in cell division in several *Nannochloropsis* species and used a luciferase reporter based system to characterize the circadian oscillations in the *Nannochloropsis* genus. In spite of the damping of cell division and transcriptional rhythms under free running conditions, the *Nannochloropsis* circadian clock displays key characteristics of a biological oscillator (Roenneberg *et al.*, 2005). This oscillator can be entrained by light and temperature, causes jet lag, leads to changes in the phase angle when entrained at periods slightly different than 24 h and leads to frequency demultiplication when the entraining period is much shorter than 24 h.

Under light/dark cycles mitosis occurs during the dark period in *N. oceanica* (Poliner *et al.*, 2015)(Figure 1) and we observed a similar phase of cell division in the four *Nannochloropsis* strains tested. Although under constant light conditions cell division rhythms were often weak, cell division occurred during the subjective night period. Similar phases of cell division under light/dark and constant light conditions have been observed in many unicellular and multicellular eukaryotic algae (Anderson *et al.*, 1985, Goto and Johnson 1995, Johnson 2010, Makarov *et al.*, 1995, Moulager *et al.*, 2007). It has been proposed that this particular phase of cell division is due to the increased UV sensitivity of the cells during the nuclear division process (Nikaido and Johnson 2000). It has also been proposed that the temporal separation of cell division from photosynthesis protects cells from excessive oxidative stress (Miyagishima *et al.*, 2014).

Bioluminescence reporters have been used widely for analysis of circadian transcriptional rhythms in a large number of species (Welsh *et al.*, 2005, Welsh and Kay 2005). We have tested several luciferase reporter genes in two *Nannochloropsis* species and shown that a firefly luciferase gene is able to track *in vivo* RNA oscillations and allows for long-term monitoring of transcription. In contrast, the *in vivo* luminescence of the small synthetic nanoluciferase tracked protein levels well but was unsuitable for long term *in vivo* monitoring. We used firefly luciferase expressing lines to analyze the fundamental properties of circadian oscillators in *Nannochloropsis*.

In contrast to the strong sustained rhythms in other unicellular algae (Noordally and Millar 2015), cell division and bioluminescence rhythms in *Nannochloropsis* species damped within 3-4 days under constant light conditions. The rhythms of *N. salina* CCMP537 appear slightly more robust than the ones from *N. oceanica* CCMP1779. The period length estimations of the bioluminescence rhythms had high variability, which might be caused by the rapid damping. In *N. salina* these rhythms appeared to be temperature compensated and had average periods of ~24-31 h. The damping of these rhythms was observed when using different promoters indicating that the decrease in amplitude was not promoter specific, although it is possible that other genes show more sustained oscillations. Future studies analyzing the genome wide transcriptome oscillations will allow us to determine the extent of circadian regulation of RNA levels in *Nannochloropsis* species. It is possible that constant light conditions inhibit transcriptional rhythms in this genus. Constant light can lead to arrhythmicity in other organisms. Circadian rhythms in the green algae *Ostreococcus tauri* are abolished under constant medium to high light intensities (Moulager *et al.*, 2010), locomotor rhythms in animals and conidiation rhythms in *Neurospora crassa* are absent under constant light conditions (Pittendrigh and Daan 1976, Schneider *et al.*, 2009), and dim light inhibits the rhythms of fruit flies (Winfree 1974). However, in *Nannochloropsis* species, we observed a similar degree of damping under high frequency T6 cycles (3 h light/3h dark) and under constant light. After one day in T6 the luminescence reporters started to react to the imposed light/dark cycles and this loss of gating correlated with the damping of the ~24 h rhythms in constant light (Figure 5), indicating that masking is not the cause of the damping of free running rhythms.

There are several potential explanations for the poor free running rhythmicity in *Nannochloropsis* species. The observed damping might be caused by desynchronization of the single cells in the culture as is the case of mammalian fibroblasts (Welsh *et al.*, 2004). It is also possible that the experimental conditions, including media composition, are not optimal for rhythmicity in the species studied (Doherty and Kay 2010, Hurley *et al.*, 2014, Olivares-Yanez *et al.*, 2016). The transcriptional rhythms in *Nannochloropsis* species could also be controlled by a damped oscillator. The dinoflagelate *Lingulodium*, a model of early circadian studies due to its robust endogenous bioluminescece rhythms, does not appear to have strong RNA oscillations under constant light conditions (Roy *et al.*, 2014) although it does display rhythms in protein synthesis (Morse *et al.*, 1989, Morse *et al.*, 1990). Future studies will reveal the role of posttranscriptional regulation of the circadian rhythms in the *Nannochloropsis* genus.

In organisms with robust circadian clocks, these clocks provide a fitness advantage (Dodd *et al.*, 2005, Woelfle *et al.*, 2004, Yerushalmi *et al.*, 2011). However, it remains unclear whether this selective advantage requires a robust self-sustained circadian oscillator or whether weaker oscillators can play a similar role. There are several photosynthetic organisms with no apparent free running oscillations. Under constant light conditions no oscillations of gene expression and chlorophyll fluorescence have been detected in the gymnosperm Norway Spruce (*Picea abies*) in spite of containing the canonical plant clock genes (Gyllenstrand *et al.*, 2014). In contrast to the widely studied *Synechococcus elongatus*, other cyanobacteria such as the abundant *Prochlorococcus* genus lack a functional *kaiA* gene and no rhythms have been detected under constant light conditions (Holtzendorff *et al.*, 2008, Schmelling *et al.*, 2017). It has been hypothesized that an hour-glass oscillator that is reset every day is sufficient for these *Prochlorococcus* species because they originate from southern latitudes where they do not experience large changes in daylength, and their marine environment is more stable than fresh water habitats (Mullineaux and Stanewsky 2009). A recent study shows that diel rhythms are stronger in *Prochlorococcus* in co-culture with a heterotrophic bacterium suggesting that biotic interactions can help maintain oscillations in plankton communities (Biller *et al.*, 2018). Experiments using arrhythmic and weakly rhythmic mutants in *S. elongatus* indicate that lines with a damped oscillator grow better than the wild type under light/dark cycles (Woelfle *et al.*, 2004) which suggests that a robust clock is not required for optimal growth under diel conditions. Non-motile phytoplankton are likely to be exposed to low light or dark conditions due to water circulation and turbulence. A weak damped oscillator might be faster to reset than a robust oscillator allowing for faster entrainment (Bordyugov *et al.*, 2015) and might be of advantage in non-motile marine plankton.

*Nannochloropsis* bioluminescence rhythms can be entrained by light and temperature cycles. The similarity between the blue and white light phase response curves indicate that a blue light receptor might mediate the main light input signal to the clock. *N. salina* phase response curves using blue light are similar to the ones observed in the green marine algae *Ostreococcus tauri* under blue light, and both display stronger advances than delays (Thommen *et al.*, 2015). In contrast to diatoms, *Nannochloropsis* species lack phytochromes (Vieler *et al.*, 2012) and it is unclear whether they contain any red light photoreceptors. *N. oceanica* encodes for three Aureochrome genes (Vieler *et al.*, 2012). Aureochromes are stramenopile specific photoreceptors that have been shown to be involved in blue light mediated photomorphogenesis in the *Vaucheria* genus and in the induction of cell division by blue light in diatoms (Kroth *et al.*, 2017, Takahashi *et al.*, 2007). Marine waters are enriched in blue light since light of longer wave lengths cannot penetrate water efficiently (Kirk 2011, Ragni and Ribera D’Alcalà 2004) and therefore blue light is likely to play a key role in signaling in marine organisms. Aureochromes contain conserved bZIP DNA-binding and LOV light-sensing motifs (Takahashi *et al.*, 2007), dimerize, and their DNA binding affinity is modulated by light exposure (Banerjee *et al.*, 2016, Heintz and Schlichting 2016, Herman *et al.*, 2013, Hisatomi *et al.*, 2013). Future studies will define the role of Aureochromes in the light input to the clock in stramenopiles.

No circadian clock components have been identified in the genus *Nannochloropsis*. *Nannochloropsis* species appear to lack clear homologs to plant clock components, although CCT domain containing proteins are interesting potential regulatory candidates. As other stramenopiles, Nannochloropsis species also encode for several proteins similar to clock components found in animals. *Nannochloropsis* CPF1 like proteins, as diatom CPF1, show similarities to 6-4 photoylases in plants and animals as well as to cryptochromes proteins involved in circadian regulation in insects and mammals (Figure S11). CPF1 from *Phaeodactylum tricornutum* as well as CPF1 from the green alga *Ostreococcus tauri* are able to inhibit the transcriptional activity of the mouse BMAL-CLOCK complex in a heterologous system, the same activity as the mouse circadian clock component CRY1 (Coesel *et al.*, 2009, Heijde *et al.*, 2010). However, although *O. tauri* does not encode for bHLH-PAS proteins, the presence of bHLH-PAS proteins in stramenopiles suggest that these interactions might be functionally relevant in stramenopiles.

## Conclusion

Using bioluminescence transcriptional reporters we provide evidence for the presence of circadian rhythms in *Nannochloropsis* species. The *N. salina* luminescence strains characterized in this study represent a powerful system for future studies to identify the mechanisms regulating circadian rhythms in stramenopiles. In addition to the luminescence reporters described in this study, the recent development of a comprehensive set of molecular tools for *Nannochloropsis* species (Poliner *et al.*, 2018a) will allow the characterization of the circadian function of the putative clock genes, the investigation of the role of photoreceptors in entrainment, and the development of mutant screens to identify lines with compromised rhythms. Future studies on the clock in stramenopiles will not only improve our understanding of the role of circadian clocks in the biology of marine phytoplankton but also provide new insight into the evolution and design principles of circadian oscillators.

## Experimental procedures

### *Nannochloropsis* species and growth conditions

The *Nannochloropsis* lines *N. oceanica* CCMP1779, *N. salina* CCMP537, *N. salina* CCMP1778, and *N. salina* CCMP1776, were originally collected at different latitudes (29°N, 41.6°N, 55.75°N, and 55.75°N respectively) as described at Bigelow National Center for Marine Algae and Microbiota (Fig. S1). We used the genome sequences and annotation versions *Nannochloropsis oceanica* CCMP1779 v1.0 (Vieler *et al.*, 2012) available at JGI and *Nannochloropsis salina* CCMP537 (Wang *et al.*, 2014) available at http://www.bioenergychina.org:8989/index.html. Liquid cultures were grown in flasks in f/2 medium and maintained on a shaker set at 120 rpm. If not otherwise indicated cultures were grown under white fluorescent lights. Blue and red light treatments were conducted using LED lights; their spectrum is shown in Figure S14.

### Growth curves

*Nannochloropsis* species were maintained in flasks with f/2 media under 100 μmol m^−2^ s^−1^ constant white light and entrained under diel conditions (12 h light/12 h dark, at 40 or 100 μmol m^−2^ s^−1^) for at least 10 days. Each culture was then diluted to make 3 50-mL cultures at the same cell density and these diluted cultures were grown in the same diel conditions for 24 h before cell counting began. Cells were counted using a Beckman-Coulter Z2 particle counter.

### Generation of luciferase expressing lines

To generate the pNOC-LUC vector series (Figure S15, Dataset 1) the Gateway-luciferase cassette from pMDC-LUC+HA (Farre and Kay 2007) and the hygromycin resistance cassette from pSELECT100 (Vieler *et al.*, 2012) were PCR amplified and combined by Gibson Assembly. The LDSP terminator (CCMP1779_4188) was PCR amplified from *N. oceanica* CCMP1779 genome using the primers LDSP term sac F+ and LDSP term afl R-(Figure S16), and digested with SacI and AflII to replace the 35S terminator as the terminator for the luciferase gene. The LUC+HA gene was then replaced by either *N. oceanica* CCMP1779 codon optimized coding sequences for firefly luciferase (FLUC) and, nanoluciferase (NLUC) (Hall *et al.*, 2012, Poliner *et al.*, 2018b) or a Chlamydomonas codon optimized renilla luciferase (RLUC) (Ferrante *et al.*, 2008) to form pNOC-Fluc, pNOC-Nluc, and pNOC-Rluc respectively using AscI and SacI and the primers listed in Figure S16. The pNOC vectors were used to transform *N. oceanica* CCMP1779. Since *N. salina* CCMP537 is resistant to hygromycin, the hygromycin resistance cassette from the pNOC-GW vectors was replaced by a zeocin resistance cassette (BleC). This cassette, which includes the CCMP537 EF promoter (NODE_7115_length_93819_cov_37.515949:78382..79716), the zeocin resistance gene and the CCMP1779 LDSP terminator, from *N. salina* selection construct pNSA-511 were cloned by restriction digest (AhdI/SbfI) generating the pNSA-FLUC and vector (Dataset 1). The pNSA-511 construct was generated by cloning the CCMP537 EF promoter into pNOC-411 using SnabI and XhoI (Poliner *et al.*, 2018b). The promoters of the cellulose synthase (NannoCCMP1779|5780, NODE_11401_length_57650_cov_57.237450:4263..6099), *LHC1* (NODE_14101_length_22726_cov_35.737923:17540..18097), and *LHC8* (NannoCCMP1779|6809, NODE_11394_length_43512_cov_40.897591:1551..2663) promoters were amplified from the *N. oceanica* CCMP1779 and *N. salina* CCMP537 genomes using primers listed in Figure S16 and inserted into pENTR-D-Topo (Invitrogen). The promoters were transferred into luciferase reporter destination vectors by an LR clonase reaction (Invitrogen) and used for transformation.

### *Nannochloropsis* transformation

Promoter reporter vectors were linearized by restriction digest, concentrated by ethanol precipitation, and resuspended in water. Transformations were conducted as described previously (Poliner *et al.*, 2018c).

### Immunoblotting

Reporter line 50 ml cultures in f/2 medium were grown in 250 ml flasks under 12 h /12 h light/dark cycles, 200 μmol m^−2^ s^−1^. Every 3 h, 5 ml aliquots were collected by centrifugation at 4,000 × G for 5 min, after decanting, pellets were transferred to 2 ml round bottom tubes and centrifuged at 13,000 × G for 30 seconds. After aspiration of the supernatant, pellets were frozen in liquid nitrogen. Frozen pellets were ground with 3 × 2 mm steel balls at 30 Hz for 2 minutes, and resuspended in protein extraction buffer (100mM Tris pH 8.0, 2mM PMSF, 2% β-mercaptoethanol, 4% SDS), heated at 80°C for 3 minutes, and centrifuged at 13,000 × G for 30 seconds. The protein concentration was determined using a RCDC protein quantification kit (Bio-rad) and equal quantities (50 μg) of protein were loaded chronologically for the light/dark timecourse, separated through 3-10% SDS-PAGE and transferred to PVDF membranes. The FLUC-HA and NLUC-HA proteins were detected with anti HA-HRP antibody (Roche 3F10) (1:1000) in TBST 5% milk. RLUC was detected with a primary anti-RLUC (MBL Life Sciences PM047) 1:2000 in TBST 5% BSA and a secondary anti-Rabbit IgG-HRP (Bio-Rad 170-6515; 1:10000) antibody in TBST 5% milk. FLUC and RLUC immunoblots were visualized with Clarity (Bio-rad) and NLUC visualized with Femto (Thermo Scientific) chemiluminescence solutions. After visualization immunoblots were stained with Direct Blue 71 protein stain (Hee-Youn Hong 2000).

### Luminescence assays

*Nannochloropsis* luciferase expressing lines were maintained and entrained under 12 h/12 h light/dark cycles (200 μmol m^−2^ s^−1^) if not otherwise indicated in the figure legends. Luminescence assays were set-up in 96-well plates with 1.5 million cells in 200 μl and 500 μM firefly luciferin (GoldBio), 100X dilution of renilla luciferin (Promega, E2920 Dual-Glo^®^ Stop & Glo^®^ Reagent), or 10,000X dilution of NanoLuc substrate (Promega, N1110) in f/2 medium per well. Luminescence assays were conducted with a Centro XS3 LB960 luminometer (Berthold Technologies) over a 0.3 sec exposure and measurements were collected every hour using MikroLab software. Bioluminescence experiments include independent cultures from 2-6 (*N. salina* CCMP537) or 3-7 (*N. oceanica* CCMP1779) independent transgenic lines (as indicated in the figure legend), with the exception of the phase response curve for which data from cultures from one transgenic *N. salina* CCMP537 *CS::FLUC* line is shown.

The casein kinase inhibitor PF-670462 (Cayman Chemicals) was resuspended in water at 2 mg ml^−1^. Pharmacological treatments were prepared using the protocol described above with some modifications. Replicate wells with equal cell quantities for each treatment and mock were prepared. Dilutions of the drug at twice the final intended concentration were prepared in the luciferin f/2 solution and 100 μl added to the respective wells containing 100 μl of cells in f/2.

For substrate refeeding experiments cultures were entrained and maintained under light/dark cycles. On the day of the experiment duplicate wells with equal cell densities and luciferin were prepared. Luminescence measurements were collected every hour. After 48 h in order to restore the starting concentration of substrate, 20 μl of f/2 and 10X concentrated substrate (10 mM firefly luciferin, 1,000X dilution of NanoLuc Dual-Glo reagent, Promega) were added to the respective wells.

### Statistical analyses

Rhythmicity was quantified using tools on Biodare 2 (https://biodare2.ed.ac.uk/)(Moore *et al.*, 2014). If not otherwise indicated in the figure legend, the raw bioluminescence traces were first detrended by subtracting a +12/-12 h moving average. To correct for damping effects, each of the resulted data points was divided by the standard deviation of the corresponding +12/-12 h moving window. This procedure is similar as one previously reported for damping circadian oscillations (Izumo *et al.*, 2006). Period length was estimated using FFT-NLS on the detrended data (Plautz *et al.*, 1997). We defined a trace as rhythmic with a circadian type period when the period length estimated with FFT-NLS was within 3 h of the estimate provided by either Enright Periodogram (ERP)(Enright 1965) or Maximum Entropy Spectral Analysis (MESA) (Burg 1972), and the FFT-NLLS Relative Amplitude Error was < 0.7, the Period Error < 2 and the Goodness Of Fit < 0.8. A good fit was further confirmed by visual inspection of the trace. For T6 experiments, traces periods of ~ 6h were considered “not rhythmic”, since there was not apparent circadian component. Other statistical analyses were implemented in GraphPad Prism.

### Phylogenetic analysis

Protein sequences derived from the *N. oceanica* CCMP1779 and *N. salina* gene models and related protein sequences were used to generate a multiple sequence alignment using the Molecular Evolutionary Genetics Analysis 7 (MEGA7) program (Kumar *et al.*, 2016) and the Multiple Sequence Comparison by Log-Expectation (MUSCLE) algorithm (Edgar 2004). The alignment file was used to create a neighbor-joining method phylogenetic tree with 1000 rounds of bootstrapping.

## Acknowledgments

We thank Hideki Takahashi for the anti-Renilla antibody, Christoph Benning for the access to the cell counter, and Stephanie Taylor for discussions and advice on the analyses of cyclic expression. We also thank Traverse Cottrell for technical assistance. This work was funded by a grant from the National Science Foundation (IOS-1354721) to E.M.F. The authors have no conflict of interest to declare.

## Short legends for supporting information

**Figure S1.** Collection location of the *Nannochloropsis* strains used in this study.

**Figure S2.** Cell division in *Nannochloropsis* species under light/dark and constant light conditions.

**Figure S3.** Effect of refeeding luciferase substrates on *in vivo* luminescence in *N. oceanica*

CCMP1779.

**Figure S4.** Overview of the period estimation analyses using Biodare2.

**Figure S5**. *In vivo* bioluminescence of *CS::FLUC* expressing lines in constant dark.

**Figure S6.** Rhythms of *N. oceanica CS::FLUC* expression under constant light under different temperatures

**Figure S7**. Bioluminescence rhythms under T6 cycles.

**Figure S8.** Phase response curve of *CS::FLUC* rhythms *in N. salina* under blue or red light.

**Figure S9.** CCT protein domains across taxa.

**Figure S10**. Expression of *N. oceanica* CCMP1779 CCT and bHLH-PAS genes under diel cycles.

**Figure S11.** Phylogenetic analysis of *N. oceanica* (CCMP1779) and *N. salina* (CCMP537) cryptochrome/photolyase proteins.

**Figure S12.** Comparison of bHLH-PAS domains across taxa.

**Figure S13**. Modulation of *CS::FLUC* rhythms by a CK1ε/δ inhibitor.

**Figure S14.** Spectrum of red and blue light sources.

**Figure S15.** Graphical representation of luciferase reporter vectors for *Nannochloropsis* species.

**Figure S16**. Primers used in this study.

**Dataset 1.** Plasmid sequences of constructs used in this study.

