## supporting information for "Identification of circadian rhythms in *Nannochloropsis* species using bioluminescence reporter lines"

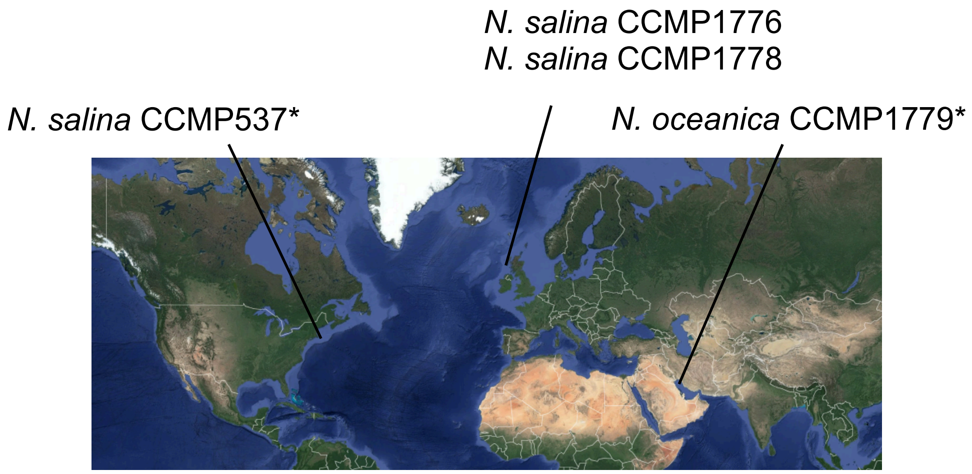

**Figure S1.** Collection location of the *Nannochloropsis* strains used in this study. Locations provided by Bigelow National Center for Marine Algae and Microbiota (ncma.bigelow.org). (*) Indicates sequenced strain. Source of image: Google Maps.

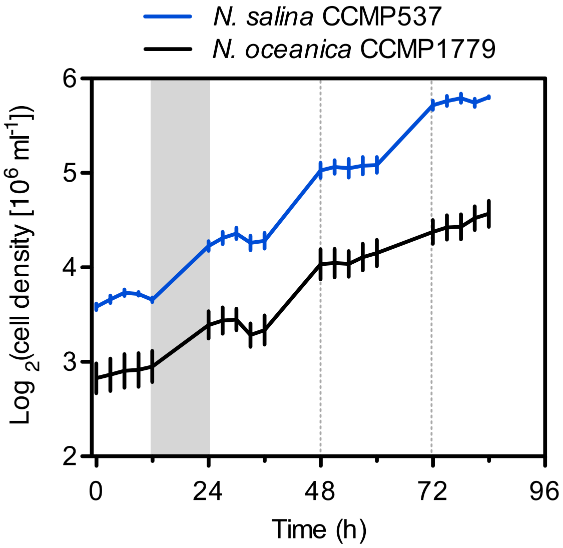

**Figure S2.** Cell division in *Nannochloropsis* species under light/dark and constant light conditions. Cultures were entrained under diel conditions (12 h light/12 h dark, 100 µmol $m^{-2}$ $s^{-1}$) for 10 days and then moved to constant light (100 µmol $m^{-2}$ $s^{-1}$). Values are the average ± SEM (n=3 cultures). Grey shading indicates dark period.

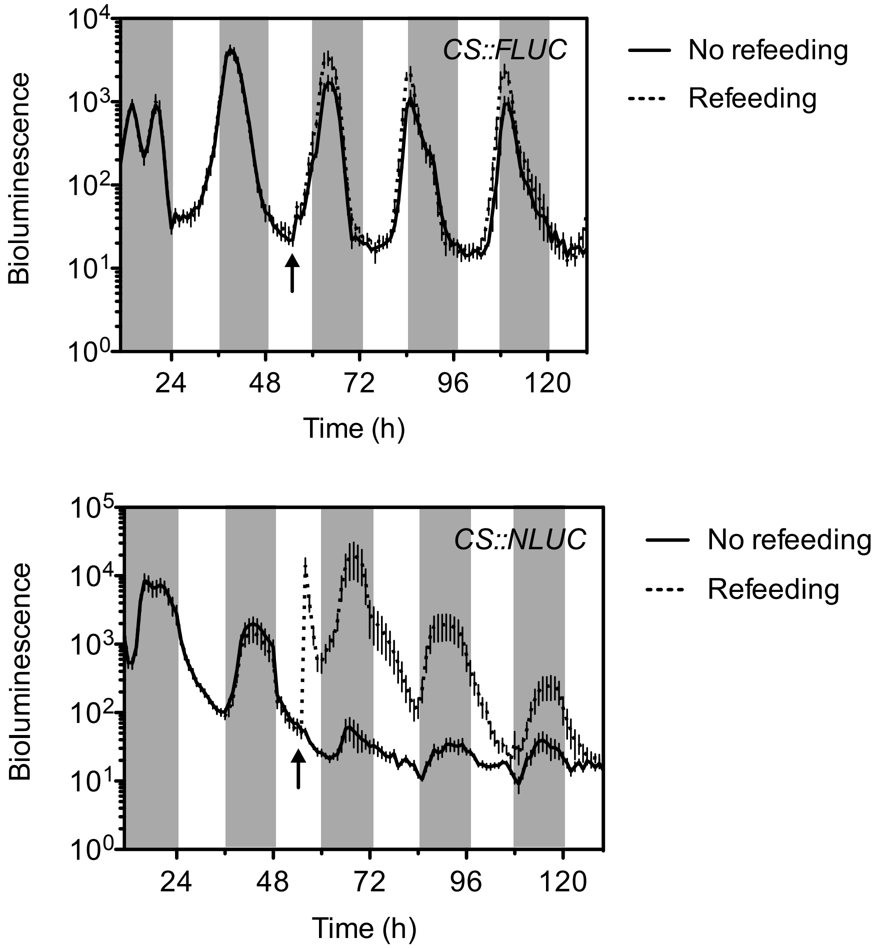

**Figure S3.** Effect of refeeding luciferase substrates on *in vivo* luminescence in *N. oceanica* CCMP1779. *In vivo* luminescence oscillations cells expressing *CS::FLUC* or *CS::NLUC* bioluminescence reporters under light/dark cycles. Dark grey shading indicates dark period. Luminescence was recorded every hour and the average bioluminescence ± SEM is shown (n=6-7 cultures, from 6-7 independent transgenic lines). The arrow indicates the time of refeeding/mock treatment. *CS*, *cellulose synthase* promoter; *NLUC*, *nanoluciferase*; *FLUC*, *firefly luciferase*.

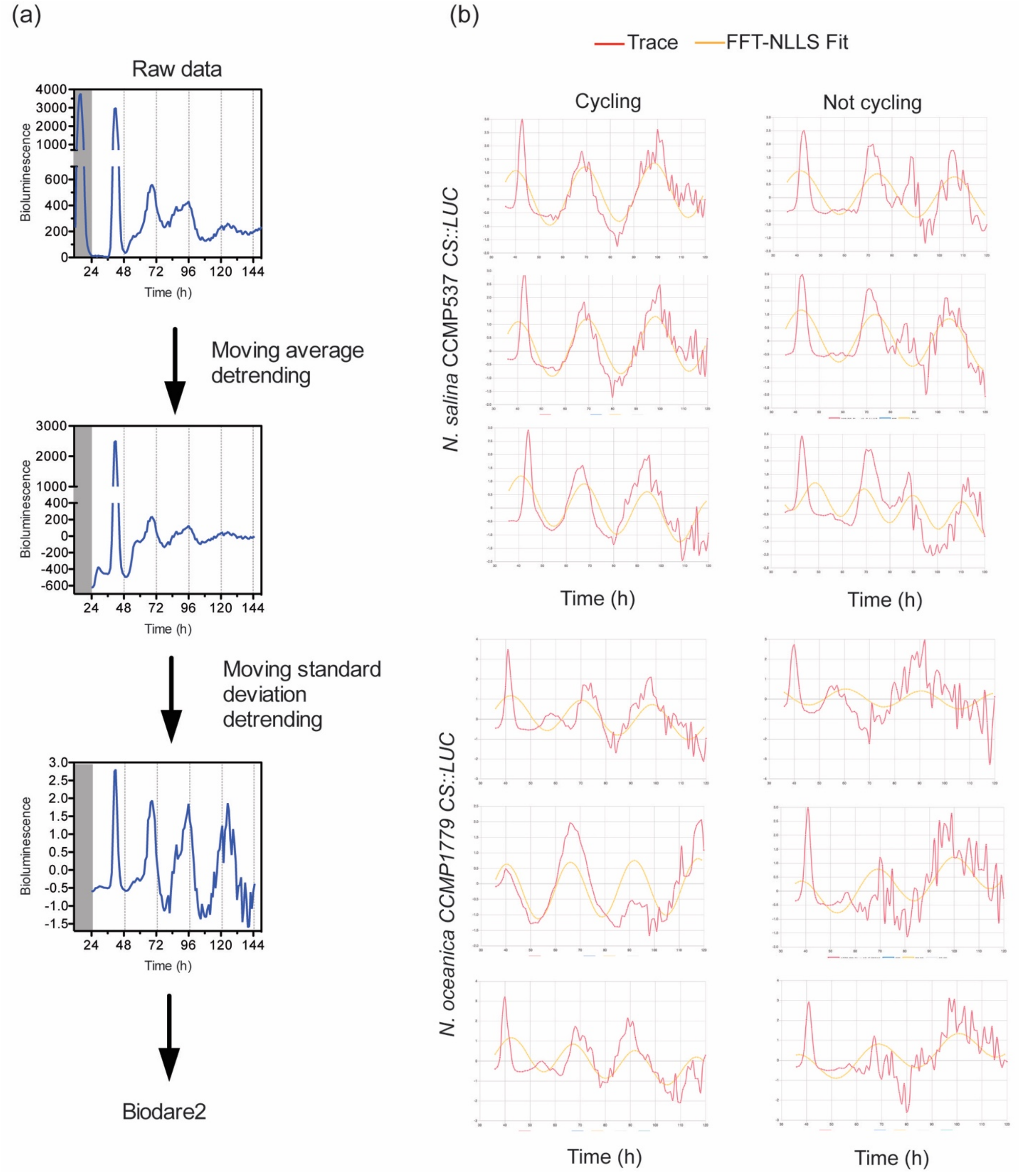

**Figure S4.** Overview of the period estimation analyses using Biodare2. **(a)** Schematic representation of the data detrending method. One trace of *N. salina CS::FLUC* under 22°C and 10 µmol $m^{-2}$ $s^{-1}$ is shown as example. Raw data were baseline detrended by substracting a moving average +/-12 h for each time point. These values were detrended for amplitude changes by dividing by a moving standard deviation (+/-12 h ) and then used for period length determination using Biodare2. **(b)** Examples of detrended data (red) and fitted curves (orange, FFT-NLLS) for traces classified as "cycling" or "not cycling" using Biodare2. Traces classified as cycling had a Period Error <2, Goodness of Fit < 0.8 and a Relative Amplitude Error < 0.7 using FFT-NLLS; the FFT-NLLS determined period should also be within +/- 3 h from the period determined by either ERP or MESA.

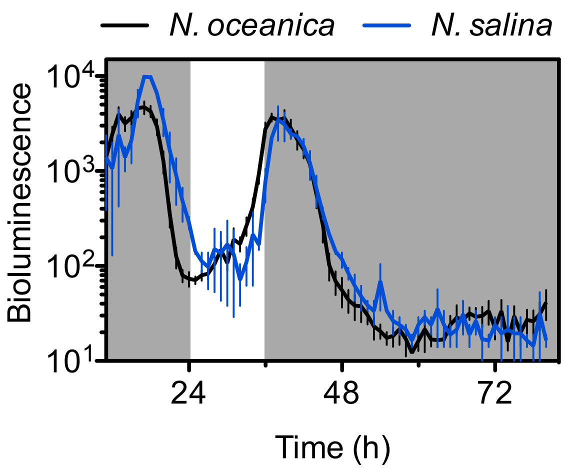

**Figure S5**. *In vivo* bioluminescence of *CS::FLUC* expressing lines in constant dark. Cultures were entrained under cycles of 12 h light/12 h dark. Dark grey shading indicates dark period. Luminescence was recorded every hour and the average bioluminescence per culture ± SEM is shown (*N. oceanica*, n=6; *N. salina*, n=2).

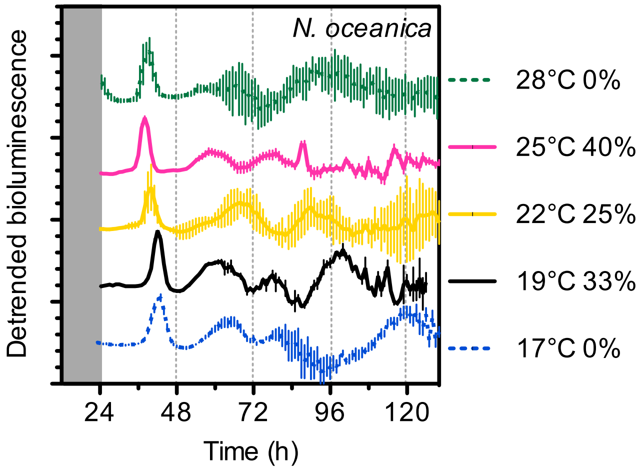

**Figure S6**. Rhythms of *N. oceanica CS::FLUC* expression under constant light under different temperatures. Cultures were entrained under cycles of 12 h light/12 h dark (200 µmol $m^{-2}$ $s^{-1}$) and 22°C, and were switched to 17°C or 19°C at time 12 h, and to 25°C or 28°C at time 24 h. The average detrended bioluminescence of rhythmic cultures ± SEM is shown (*N. oceanica* n=6-18, three independent transgenic lines), with the exception of conditions with no rhythmic traces, in which the total average is shown as a dotted line. Percent of rhythmic traces are indicated. Traces are nudged to aid visualization.

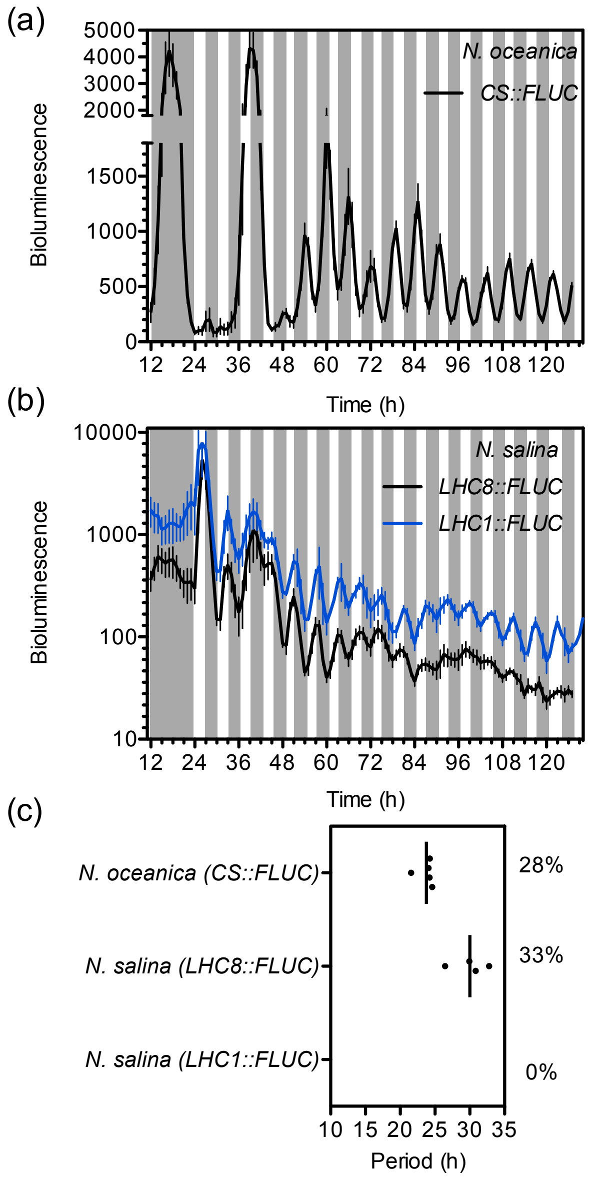

**Figure S7**. **Bioluminescence rhythms under T6 cycles**. **(a)** *In vivo* bioluminescence of *CS::FLUC* in *N. oceanica*. Grey shading indicates dark period. Cultures were entrained under cycles of 12 h light/12 h dark and 22°C (200 µmol m$^{-2}$ s$^{-1}$). Bioluminescence was measured for one night period and during cycles of 3 h white light (100 µmol m$^{-2}$ s$^{-1}$) and 3 h of dark at 22°C. The average bioluminescence of rhythmic cultures ± SEM is shown (n=5). **(b)** *In vivo* bioluminescence of *N. salina*. Values are the average bioluminescence per rhythmic culture ± SEM for *LHC8::FLUC* (n=4); and the average of all analyzed cultures for *LHC1::FLUC* (n=8)(no rhythmic traces). Cells were treated as in (a). **(c)** Period length estimated using FFT-

NLLS on Biodare2 using data shown in (a) and (b), line indicates average. Percent rhythmic traces are indicated.

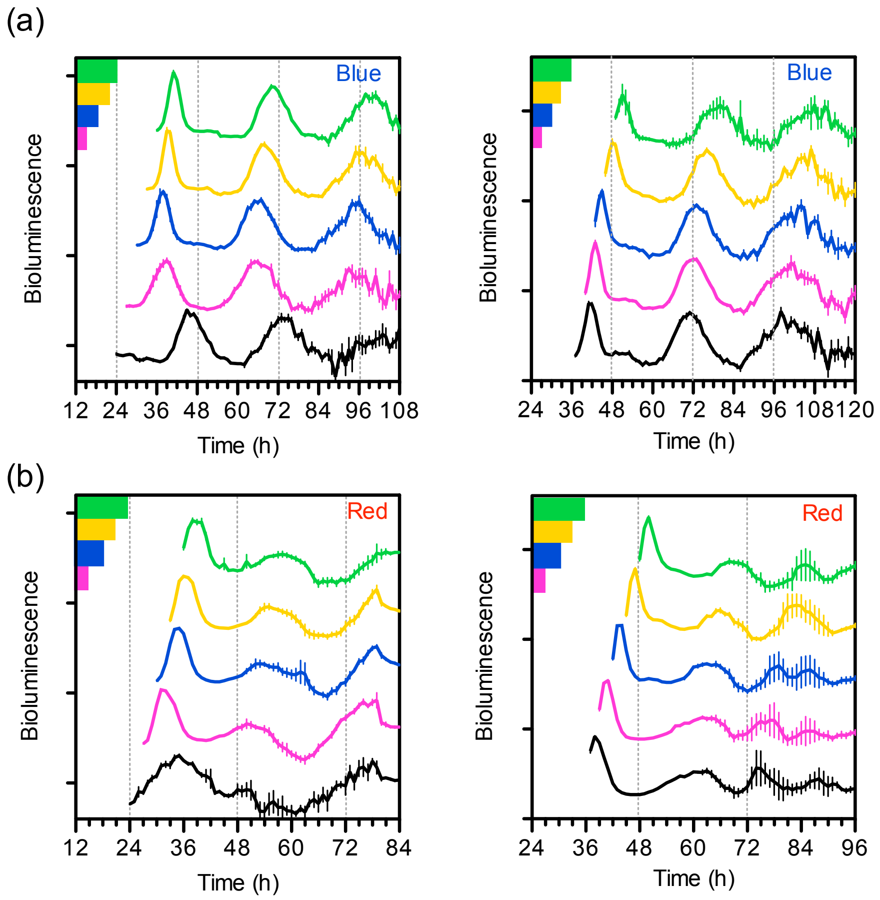

**Figure S8.** Phase response curve of *CS::FLUC* rhythms *in N. salina* under blue or red light. After the cells were entrained under cycles of 12 h light/12 h dark and 22°C for 7 days, they were exposed to one dark period of variable length before being transferred to constant blue **(a)** or red **(b)** light (10 µmol $m^{-2}$ $s^{-1}$). The detrended *in vivo* luminescence for one transgenic line (average ± SEM, n = 4) is shown. Left panels show experiments with extended nights, in which the length of the dark period was 12 h (black), 15 h (magenta), 18 h (blue), 21 h (yellow) or 24 h (green). Right panels show experiments with short nights, in which the length of the dark period was 0 h (black), 3 h (magenta), 6 h (blue), 9 h (yellow) or 12 h (green).

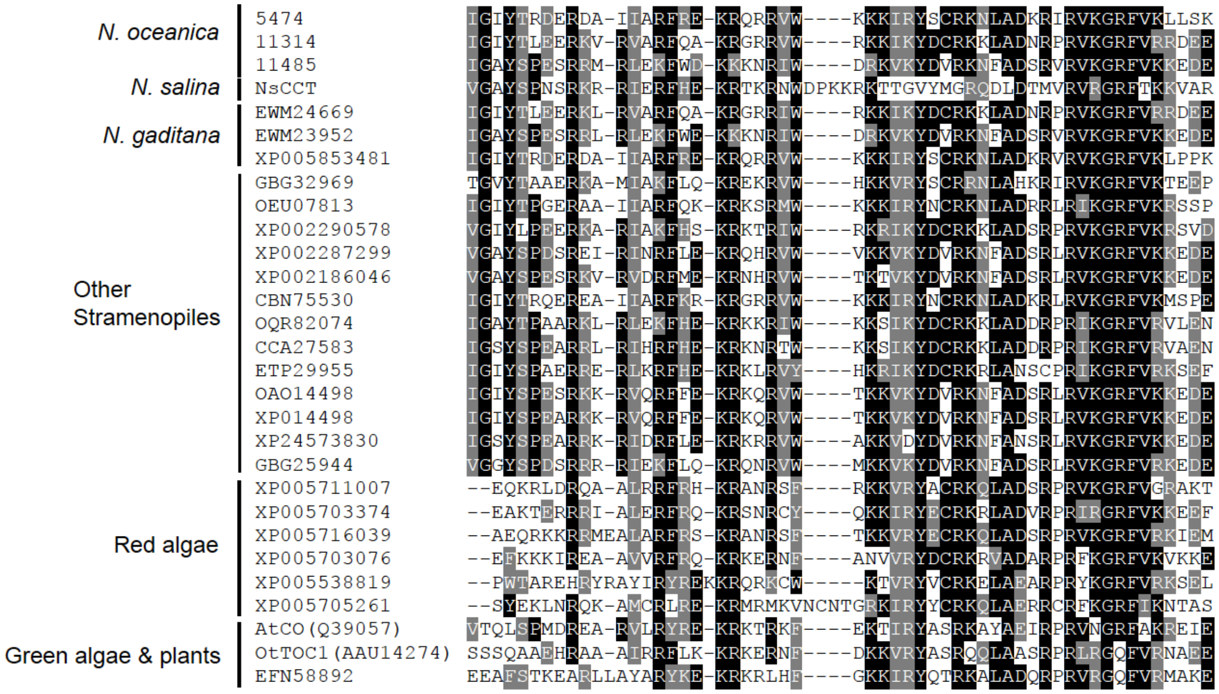

**Figure S9.** CCT protein domains across taxa. Proteins were aligned using Muscle. Shading was performed using Boxshade and amino acids identical or similar in more than 50% of the sequences are shaded. *N. oceanica* CCMP1779 numbers indicate the protein ID (CCMP1779|#) (CCMP1779 V1.0 annotation, https://genome.jgi.doe.gov)(Vieler *et al.,* 2012). The *N. salina* CCMP537 sequence (NsCCT) is evm.model.NODE_2480_length_85397_cov_42.408234.19 (Wang *et al.,* 2014). All other IDs are from Genbank. AtCO (*Arabidopsis thaliana* CONSTANS) and OtTOC1 (*Ostreococcus tauri* TOC1) have been functionally characterized (Corellou *et al.,* 2009, Yanovsky and Kay 2002).

(a)

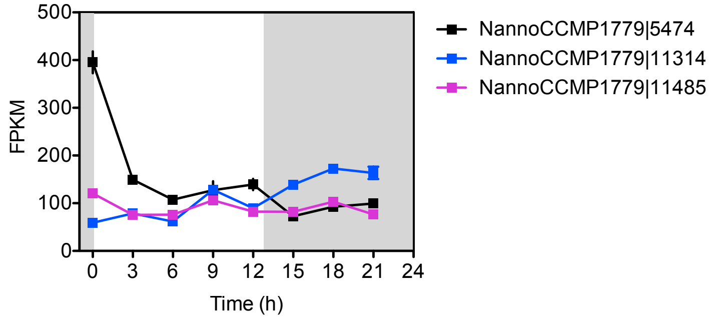

(b)

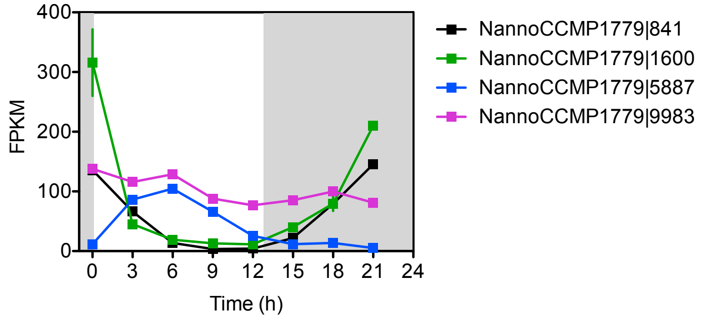

**Figure S10**. Expression of *N. oceanica* CCMP1779 CCT and bHLH-PAS genes under diel cycles. **(a)** Expression of CCT domain containing genes. **(b)** Expression of bHLH-PAS domain containing genes. Dark shading indicates dark period. Data (average ± range, n = 2) from (Poliner *et al.,* 2015).

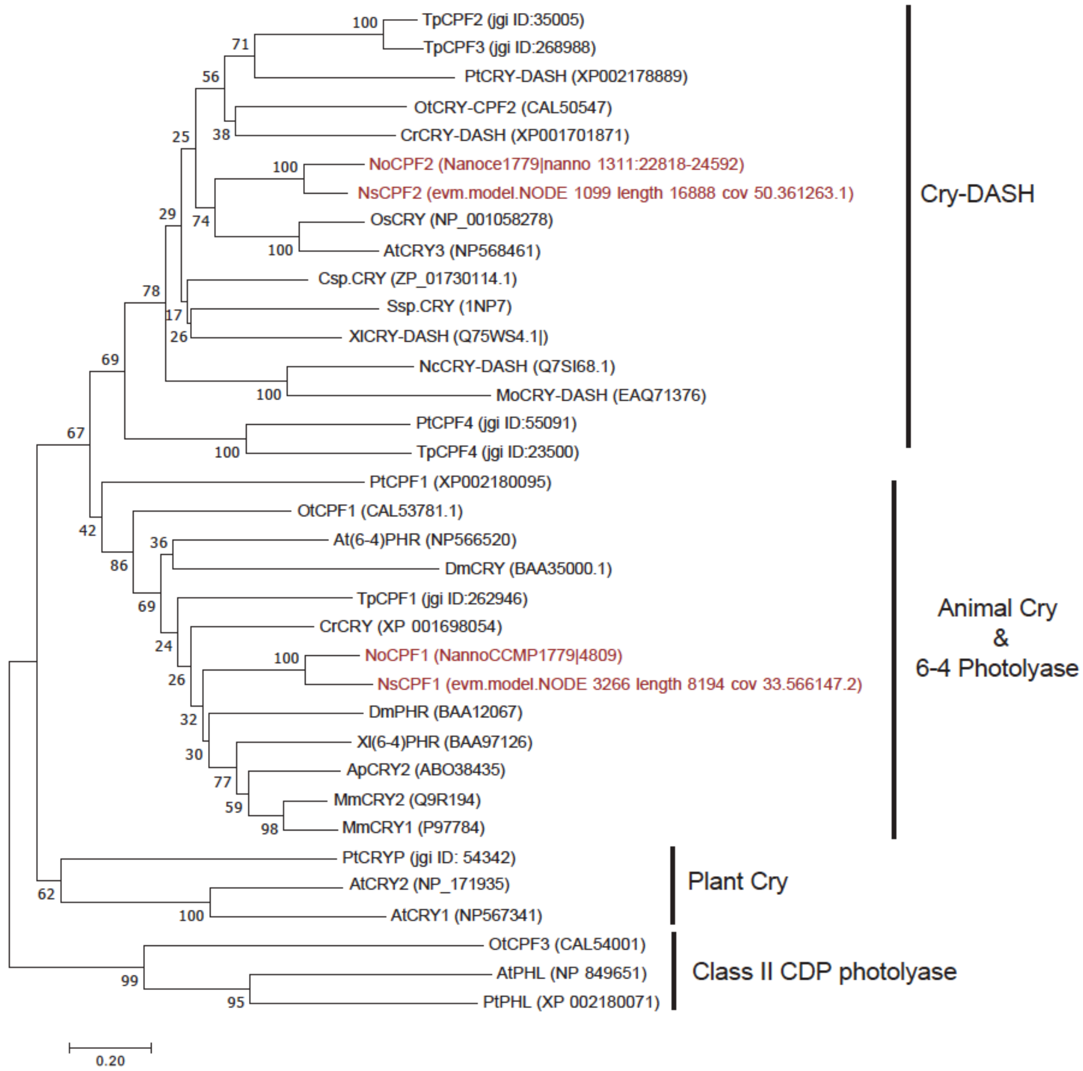

**Figure S11.** Phylogenetic analysis of *N. oceanica* (CCMP1779) and *N. salina* (CCMP537) cryptochrome/photolyase proteins. Phylogenetic analysis using the neighbor-joining method of a Muscle alignment of 35 proteins was performed in MEGA 7. The percentage of replicate trees in which the associated taxa clustered together in the bootstrap test (1000 replicates) are shown next to the branches. The evolutionary distances were computed using the JTT matrix-based method. Scale bar, 0.2 substitutions per site. NoCPF2 was not identified within the proteins annotated in the CCMP1779 V1.0 annotation of the genome, it was identified by Blast using the translated genomic sequence (CCMP1779 V1.0, https://genome.jgi.doe.gov)(Vieler *et al.,* 2012). *N. salina* CCMP537 sequence IDs are from (Wang *et al.,* 2014). The ID numbers for some diatom proteins are according to the annotated genomes at http://genome.jgi-psf.org/.

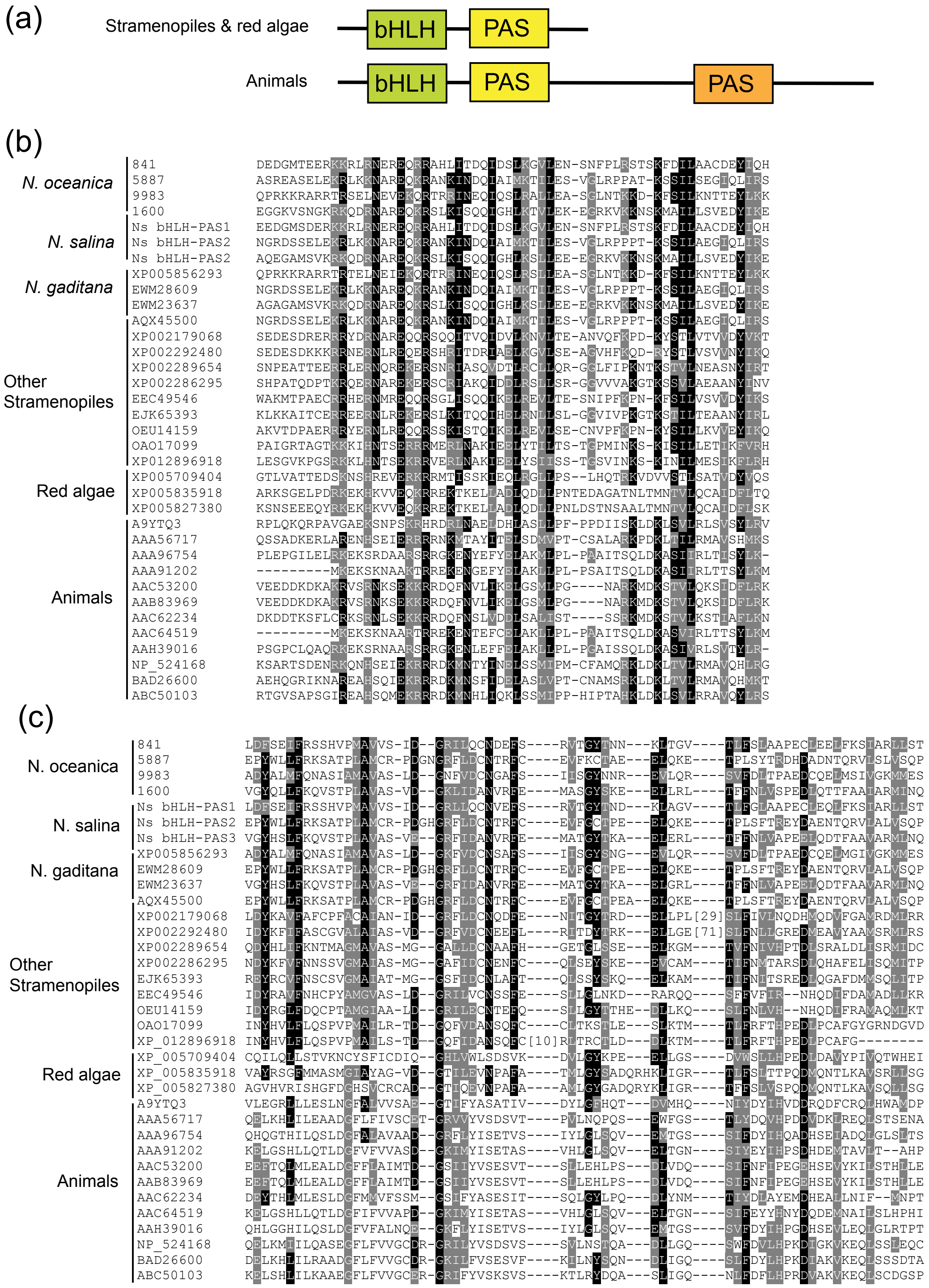

**Figure S12.** Comparison of bHLH-PAS domains across taxa. **(a)** Schematic representation of the structure of bHLH-PAS proteins in stramenopiles and animals. **(b)** Alignment of the bHLH domain of stramenopile, red algae and animal bHLH-PAS proteins. **(c)** Alignment of the PAS domain closer to the bHLH domain of stramenopile, red algae and animal bHLH-PAS proteins. For (**b)** and (**c)** proteins were aligned using Muscle. Shading was performed using Boxshade and amino acids identical or similar in more than 50% of the sequences are shaded. *N. oceanica* CCMP1779 numbers indicate the protein ID (CCMP1779|#)(CCMP1779 V1.0, https://genome.jgi.doe.gov)(Vieler *et al.,* 2012). *N. salina* CCMP537 sequence IDs are from (Wang *et al.,* 2014): Ns bHLH-PAS1, evm.model.NODE_15061_length_171711_cov_31.321459.28; Ns bHLH-PAS2, evm.model.NODE_7751_length_15947_cov_24.350849.6; Ns bHLH-PAS, evm.model.NODE_3057_length_6465_cov_23.472235.1.

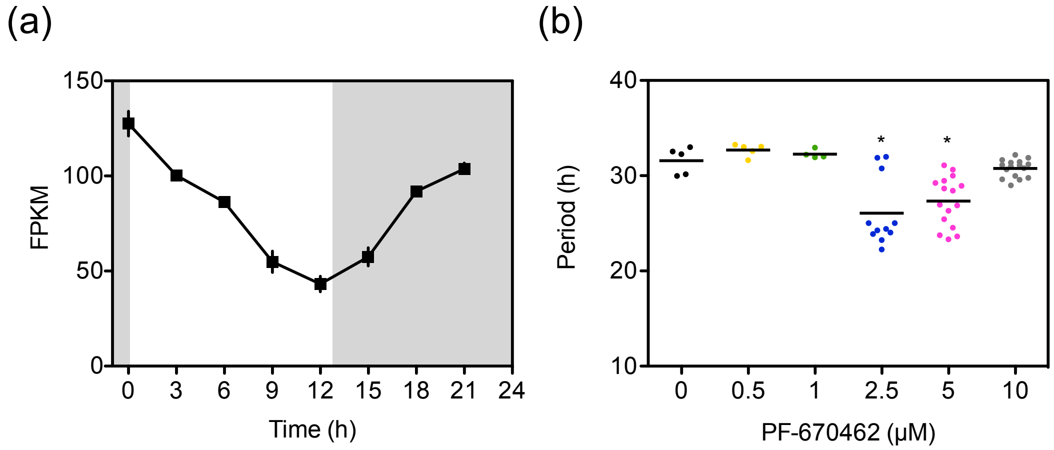

**Figure S13**. Modulation of *CS::FLUC* rhythms by a CK1ε/δ inhibitor. **(a)** Expression of a CK1 gene in *N. oceanica* (NannoCCMP1779| 10930) under light/dark cycles. Dark shading indicates dark period. Data (average ± range, n = 2) from (Poliner *et al.,* 2015). **(b)** Average period of *in vivo* bioluminescence (n=8-16) estimated using FFT-NLLS on Biodare 2. *N. salina* cells were treated with PF-670462 as described in Fig. 8. * Indicate a significant difference with the vehicle control (one-way ANOVA with Dunnett's post hoc test, α=0.05)

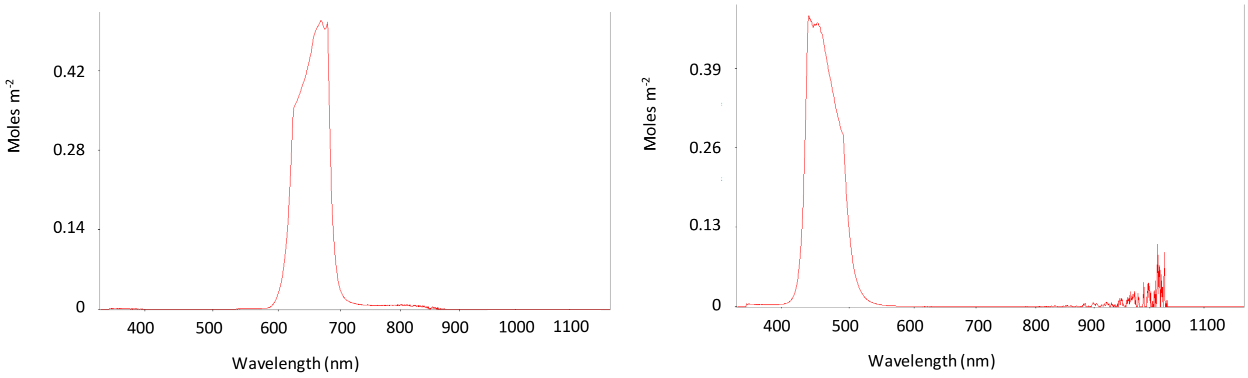

**Figure S14.** Spectrum of red and blue light sources. Spectrum was measured using a StellarNet EPP2000 VIS-50 spectrometer.

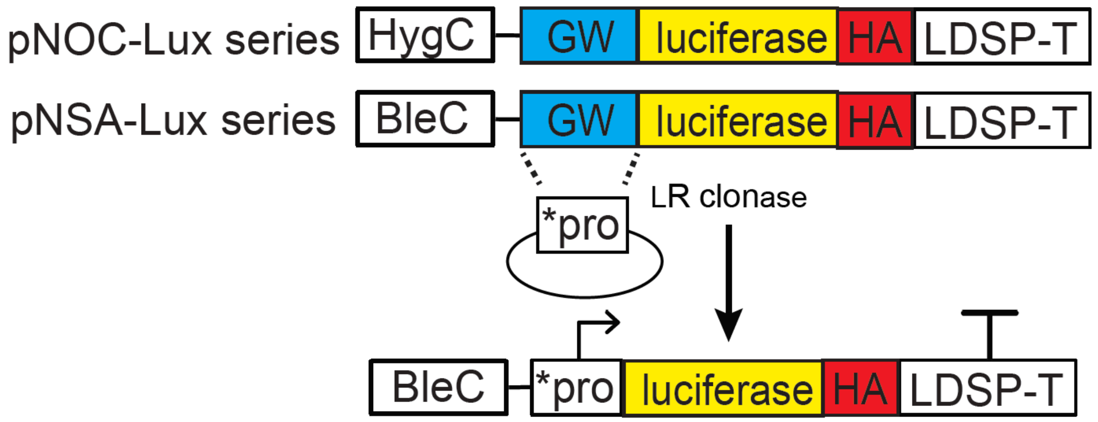

**Figure S15.** Graphical representation of luciferase reporter vectors for *Nannochloropsis* species. pNOC-LUC series are for *N. oceanica* and contain a hygromycin resistance gene regulated by the *N. oceanica* CCMP1779 LDSP promoter (HygC). pNSA-LUC series are for *N. salina* and contain a Zeocin resistance gene regulated by the *N. salina* CCMP537 EF promoter (BleC). Target promoters (*pro) can be cloned via a LR clonase reaction into the gateway cassette (GW). Each vector contains a *Nannochloropsis* codon-optimized luciferase (either firefly, Nanoluciferase, or Renilla) with a N-terminal HA tag, and the CCMP1779 LDSP terminator (LDSP-T).

| | |
|---|---|
| **Destination vector cloning** | |
| LDSP term sac F+; | GAGCTCGAAAGATCCAAGAGAGACGAG |
| LDSP term afl R-; | CTTAAGGTGATGCTGTTGCTCTTTCC |
| Renilla asci12 F2+; | GGCGCGCCTCATGGGCGGATCCGGCGCCAGCAAGGTGTACGAC |
| Renilla saci R2-; | gagctcTTACGTATCGTTCTTCAGCACG |
| Nanno Nlux asc F+; | GGCGCGCCTCATGGTGTTTACTCTCGAGGACTTCGTGGGCGACTGGC |
| Nanno Nlux sac R2-; | GAGCTC A CGCGTAGTCGGGCACGTCGTAGGGGTATCCGGCCAGGATGC |
| CCMP537 EFpro snabI F+ | caacaatacgtaCGATCCTTCTGATTGTTTGTC |
| CCMP537 EFpro xhoI R- | cagctgCTCGAGGGTTTACAGATTAGGTGTTG |
| **CCMP1779 promoters** | |
| CCMP1779 Cspro GW F+ | caccGTTTTATGTTCTATTGAGCGG |
| CCMP1779 Cspro start GW R- | CATGGTTGCGGTGTCTTACAGC |
| CCMP1779 LHC8pro GW F+ | caccGCAGCTTGCGTTTCTTTTTC |
| CCMP1779 LHC8pro GW R- | AGAGATGTGGATATTTTTGAGGTG |
| **CCMP537 promoters** | |
| CCMP537 CSpro GW F+ | cacc GTAACATCGGCAAAATTAGG |
| CCMP537 CSpro GW R- | CATGGCTGCAAAATGACTAG |
| CCMP537 LHC1pro GW F+ | CACCGTTGTTTGTGCCAAAAGCTGTC |
| CCMP537 LHC1pro GW R- | CATGAGGGAGGAGACGGTG |
| CCMP537 LHC8pro GW F+ | caccGCATATCTTCGGCTTCAGG |
| CCMP537 LHC8pro GW R- | CATGCTTCAGCTTTACAATGCG |

**Figure S16**. Primers used in this study.
